# Persistence and stability of interacting species in response to climate warming: The role of trophic structure

**DOI:** 10.1101/2020.02.28.970012

**Authors:** Taranjot Kaur, Partha Sharathi Dutta

**Affiliations:** Department of Mathematics, Indian Institute of Technology Ropar, Rupnagar, Punjab 140 001 India

**Keywords:** global warming, food web modules, omnivory, competition, species co-existence, biodiversity loss

## Abstract

Over the past century, the Earth has experienced roughly 0.4–0.8°C rise in the average temperature and which is projected to increase between 1.4–5.8°C by the year 2100. The increase in the Earth’s temperature directly influences physiological traits of individual species in ecosystems. However, the effect of these changes in community dynamics, so far, remains relatively unknown. Here we show that the consequences of warming (i.e., increase in the global mean temperature) on the interacting species persistence or extinction are correlated with their trophic complexity and community structure. In particular, we investigate different nonlinear bioenergetic tri-trophic food web modules, commonly observed in nature, in the order of increasing trophic complexity; a food chain, a diamond food web and an omnivorous interaction. We find that at low temperatures, warming can destabilize the species dynamics in the food chain as well as the diamond food web, but it has no such effect on the trophic structure that involves omnivory. In the diamond food web, our results indicate that warming does not support top-down control induced co-existence of intermediate species. However, in all the trophic structures warming can destabilize species up to a threshold temperature. Beyond the threshold temperature, warming stabilizes species dynamics at the cost of the extinction of higher trophic species. We demonstrate the robustness of our results when a few system parameters are varied together with the temperature. Overall, our study suggests that variations in the trophic complexity of simple food web modules can influence the effects of climate warming on species dynamics.

## Introduction

Increasing heat waves, severe thunderstorms, rising sea levels, coral bleaching, and loss of ecosystems are a few indicators of current unprecedented global warming (IPCC 2018). Temperature is a major abiotic factor that causes variation in species interactions and abundances (Parmesan and Yohe 2003, Deutsch et al 2008). It is well established that climate warming can lower species abundance, which in turn may affect their persistence (Parmesan and Yohe 2003). In fact, the increasing temperature creates negative impacts on populations, pushing our ecosystems closer to the face of mass extinction and a considerable loss of biodiversity (Drake and Lodge 2004, Urban 2015, Ceballos et al 2017). However, the understanding of the consequences of warming on ecological communities and individual species is still in its infancy, as warming simultaneously affects different levels of biological organizations ranging from changes in species physiological traits to their interaction networks (Fussmann et al 2014, Uszko et al 2017, Englund et al 2011).

Several experimental and theoretical studies have reported diverse impacts of warming on ecosystem stability and persistence (Vasseur and McCann 2005, O’Connor et al 2011, Binzer et al 2012, Uszko et al 2017, Rudolf and Roman 2018). For instance, Vasseur and McCann (2005), while framing a bioenergetic consumer-resource model, predicted that warming does not cause species extinctions, whereas, with a decrease in the inverse enrichment ratio, it increases the tendency to show oscillations in species abundance. In a herbivore and plant model, O’Connor et al (2011) showed that warming could result in increased stability but decreased persistence of the species. Ohlberger et al (2011) used a size-structured fish population model and predicted that warming might increase intraspecific competition among consumers, and induce regime shifts that destabilize population dynamics. The variation in the impacts of warming on ecosystems observed in recent studies emerges from the interactions among biological and physical processes as well as species associations with one another.

Species response to different biotic and abiotic factors is complex, and often induce changes in the foraging (e.g., attack rate, handling time) and life-history (e.g., growth rate, reproduction, survivorship) traits. Recent evidences show that the strength of biotic factors on population dynamics is influenced by the strength of abiotic factors between them and vice-versa (Holling 1973, Walther et al 2002, Traill et al 2010, O’Connor et al 2011, Atkinson and Urwin 2012, Post 2013, Fussmann et al 2014). On that account, several empirical studies reveal monotonically increasing relationship between species performance/biological traits with increasing temperature (Van der Have and De Jong 1996, Gillooly et al 2001, Savage et al 2004). The monotonically increasing formulations of growth rate and metabolic rates have been widely used by ecologists (Vasseur and McCann 2005, O’Connor et al 2011, Binzer et al 2012, Fussmann et al 2014, Gilbert et al 2014, Amarasekare 2015), and forms the foundation of the Metabolic Theory of Ecology (MTE) (Brown et al 2004). Yet, there are evidences that the thermal habituation of attack rates and handling times of most ectotherms represents unimodal functions of temperature (Englund et al 2011, Dell et al 2011, Amarasekare 2015); the attack rate varies in a bell-shaped manner while the handling time varies in a U-shaped fashion (Zamani et al 2006, Dell et al 2011, Englund et al 2011, Clusella-Trullas et al 2011, Amarasekare 2015). As the declining performance-temperature relationship has been rarely incorporated into ecological models studied in the recent past, understanding the consequences of warming on trophic interactions forms a major knowledge gap. Thus, it is important and timely to emphasize the interplay between species realistic biotic factors with the abiotic factors.

Majority of the recent theoretical studies on the impacts of changing temperatures on species interactions and performance have considered models of pairwise consumer-resource interactions, to maintain the mathematical tractability (Vasseur and McCann 2005, O’Connor et al 2011, Fussmann et al 2014, Gilbert et al 2014, Uszko et al 2017). However, ecological communities are composed of complex trophic interactions (McCann et al 1998, Davis et al 1998), and models restricted only to lower levels of trophic interactions can put light on limited ecological consequences. As various mechanisms that regulate co-existence or abundance of species, e.g. top-down hypothesis or green-world hypothesis (Levin 1970, Fretwell 1987, Hairston Jr and Hairston Sr 1993) cannot be considered while investigating lower-order trophic interactions. Furthermore, trophic structures of ecosystems play a significant role in propagating the immediate as well as concurrent effects of warming on species dynamics (Tylianakis et al 2008, Cardinale et al 2012, Urban et al 2017). For instance, warming can affect direct as well as indirect negative feedbacks within a species, which may suppress or fuel the oscillatory dynamics of the system (Johnson and Amarasekare 2014). Further, a rise in the temperature may disrupt the performance of an invading species (in a competitive interaction), which may moderately reduce the adverse effects of warming on other competitors (Rudolf and Roman 2018). Thus, exploring the influence of warming in population models involving complex trophic interactions can provide a broader scope to understand ecosystem behaviors (Pimm et al 1991, Murdoch et al 2003, Touboul et al 2018).

To predict the ecological responses to climate change, Davis et al (1998) and Tylianakis (2009) experimentally demonstrated the importance of understanding the interaction between temperature dependent consumer-resource systems in altering food web dynamics. Also, Edwards and Richardson (2004) studied how warming affects marine pelagic communities leading to a lack of congruence between the trophic levels. Later, Binzer et al (2012) investigated a food chain model that incorporates body mass and temperature dependencies of species traits, and projected extinction of higher trophic species. Nonetheless, we still have a limited understanding of how complex trophic interactions can alter the consequences of warming on species persistence and stability.

To fill this knowledge gap, here we use frameworks that integrate temperature dependence of species biological traits into trophic dynamics and study the influence of different community structures/food web modules in altering warming effects on species persistence and stability. By stability, here we mean the tendency of species to exhibit oscillation free behavior (i.e. system exhibits stable equilibrium forming an interior point attractor) (McCann and Hastings 1997). The food web modules have been studied in the order of their increasing complexity, where complexity is described as the product of species richness (i.e., diversity) and connectance (MacArthur 1955, May 1972, 1973, Landi et al 2018). We begin with analyzing the dynamics of a bioenergetic tri-trophic food chain model (Fretwell 1987, Binzer et al 2012) under warming. Further, we introduce an additional intermediate consumer (competitor) in the food chain, forming a diamond food web (Leibold 1996). After that, by increasing connectance of the food chain, forming an omnivorous structure, we deduce how omnivory together with warming influences the stability of the system (Pimm and Lawton 1978, Pimm 1982). Specifically, by modulating the monotonically increasing temperature dependence of species growth rates and metabolism as well as the unimodal temperature dependence of foraging rates, we focus on the consequences of warming on individual species dynamics as well as on envisioning stability and co-existence of species in different trophic structures.

Here, we find that at high temperatures, the unimodal temperature dependence of species foraging abilities along with increasing metabolism pushes higher trophic species towards extinction, irrespective of their interaction network. Further, the influence of warming on species persistence and stability varies on changing their trophic complexity. An increase in the complexity from the food chain to the omnivory reduces oscillatory dynamics in the system. The co-existence of species falls as top-down control fails in the diamond food web, and also warming does not support the intermediate consumer in the omnivorous interaction. Moreover, through sensitivity analysis, we demonstrate that our observations are valid in a rather large region in parameter space. Overall, we find that the increase in the trophic complexity can alter the impact of climate warming on species dynamics. Though, it increases ecosystem stability, yet decreases species persistence.

## Models

Here, we analyze tri-trophic food web modules (see Fig. 1) with temperature-dependent traits. These models include a resource, either one or two intermediate consumers, and a top predator. First, we consider a food chain model.

**Figure 1.**
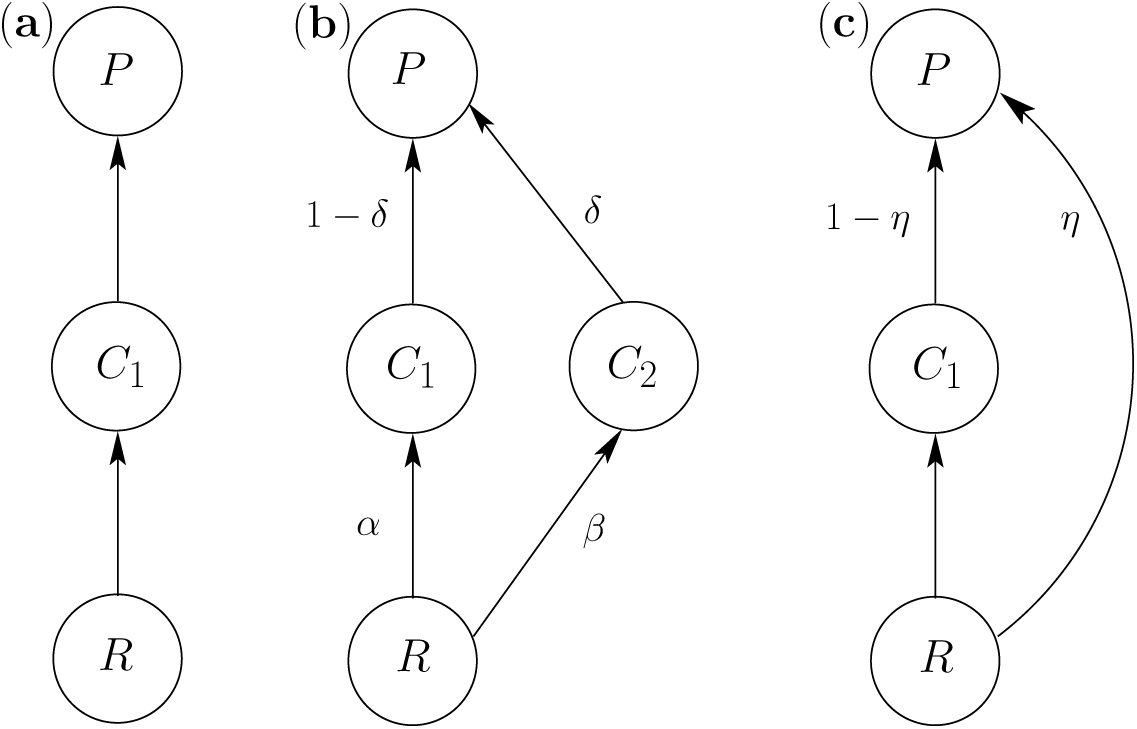
Schematic representations of the considered food web modules. (a) Interaction within a basal resource (*R*), an intermediate consumer (*C*_1_), and a top predator (*P*) in a tri-trophic food chain. (b) A diamond food web, where *α, β, δ*, and 1 − *δ* signify the interaction preferences between the linked species. (c) Interaction within the species when the predator *P* is an omnivore, where *η* determines the strength of omnivory. The arrows represent the direction of energy flow from one species to another.

### Tri-trophic food chain

A tri-trophic food chain is composed of a fundamental energy pathway between a resource, a consumer, and a top predator. We consider a basal resource with a density *R*, acquired by a consumer having density *C*_1_, which further acts as an energy source for a top predator with density *P* (see Fig. 1(a)). The food chain model has the following form:

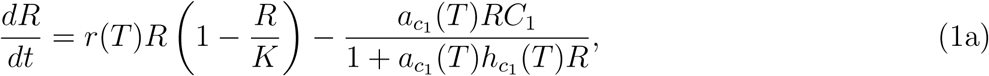

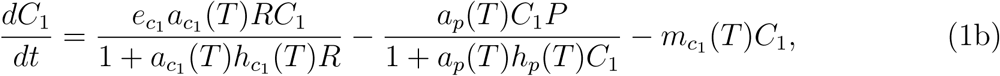

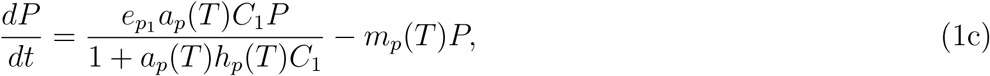

where the basal resource follows temperature-dependent logistic growth having intrinsic growth rate *r*(*T*) and carrying capacity *K*. Species consumption follows the Holling Type-II functional response (Murdoch et al 2003) which is characterized by saturating intake rate. 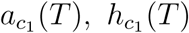 and 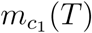 are temperature influenced attack rate, handling time and metabolism for the intermediate consumer *C*_1_, respectively. 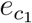 determines *C*_1_’s efficiency in absorbing and converting the acquired resource into energy, known as the conversion efficiency. The conversion efficiency is generally considered to be unaltered by temperature (Vasseur and McCann 2005, Johnson and Amarasekare 2014, Uszko et al 2017). Moreover, the stoichiometric estimates of conversion efficiency of the organisms indicate it to be insensitive to thermal constraints (Custer 2005). Similarly, *a*_*p*_(*T*), *h*_*p*_(*T*) and *m*_*p*_(*T*) are the attack rate, handling time, and metabolic rate, respectively, of the top predator. The conversion efficiency of *P* on consuming *C*_1_ is given by 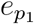

In Eqn. (1), the saturating Type-II functional response of *C*_1_ on *R* is positive feedback on resource per-capita growth rate, making the resource-consumer link oscillatory. Further, Type-II functional response of *P* on *C*_1_ can amplify the influence of environmental perturbations between the consumer-predator link. The dampening of oscillations in both the links is governed by the negative feedback on resource per-capita growth rate, explicitly due to resource self-limitation and implicitly as a consequence of predation (Johnson and Amarasekare 2014).

### Temperature dependence of species phenotypes

Using empirical evidences, many studies on the temperature dependence of consumer-resource interactions have considered unimodal or monotonically increasing thermal response of species biological traits (Vasseur and McCann 2005, Gilbert et al 2014, Amarasekare 2015, Dell et al 2014). The inclusion of temperature as one of the influential parameters in species dynamics mainly originates from two sources: First, the increase in the kinetic energy due to increase in the temperature leads to a higher collision of molecules that are responsible for various biochemical reactions to take place (Sharpe and DeMichele 1977, Schoolfield et al 1981, Kingsolver 2009, Kingsolver et al 2011). Second, the temperature response of ectothermic taxa has also been reported in studies that use large scale empirical data (Englund et al 2011, Kingsolver et al 2011, Dell et al 2011).

Here, temperature is incorporated into the metabolic rates of species (ectotherms) by using the Boltzmann-Arrhenius relationship for reaction kinetics (Van der Have and De Jong 1996, Gillooly et al 2001, Savage et al 2004, Brown et al 2004). The consumer’s and the predator’s metabolism 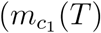 and *m*_*p*_(*T*), respectively) increases monotonically with temperature (*T*). The temperature dependence of the parameters mentioned above is formulated as:

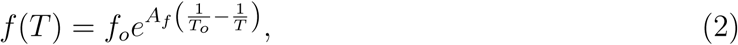

where *f* (*T*) is the species phenotype at temperature *T*, *f*_*o*_ is the trait value at the ideal (reference) temperature *T*_*o*_ (typically lying between 24°C-25°C) (Sharpe and DeMichele 1977, Schoolfield et al 1981). *A*_*f*_ corresponds to the Arrhenius constant which measures the trait sensitivity towards temperature (see supplementary information SI-1, Eqns. (SI-1.1)–(SI-1.4)). Empirical studies account for both the monotonically increasing/unimodal temperature dependence of resource growth rate. Here, we consider that resource intrinsic growth rate *r*(*T*) increases with increases in the mean temperature (Vasseur and McCann 2005, Van de Wolfshaar et al 2008, Amarasekare 2015), given by Eqn. (2). The unimodal temperature dependence of *r*(*T*) has been discussed in detail in the supplementary SI-3.

Species thermal performance norms track the performance of their phenotypes under thermal constraints (Zamani et al 2006). Thermal performance curve (TPC) of the attack rate (the handling time) increases (decreases) at low-temperature values, attain optimum value at an ideal temperature and later declines (inclines) along the temperature gradient (Deutsch et al 2008, Vasseur et al 2014). Moreover, recent studies show that foraging traits of a large number of ectotherms like lizards, snakes exhibit symmetric, unimodal temperature response (Dell et al 2011, Clusella-Trullas et al 2011, Englund et al 2011, Amarasekare 2015). Thus, temperature-dependent attack rates 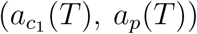 and handling times 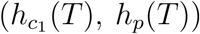 of a species explaining their TPC’s are modelled using the Gaussian function and in this case *f* (*T*) becomes (Amarasekare 2015):

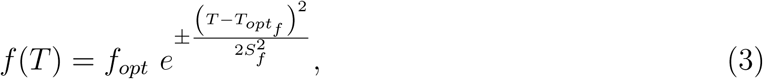

where 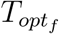 is the temperature corresponding to the optimum trait value *f*_*opt*_ and *S*_*f*_ is the performance breadth for the temperature-dependent trait *f* (*T*). It quantifies the temperature sensitivity of the corresponding biological trait of the species (see supplementary information SI-1, Eqns. (SI-1.5)–(SI-1.10)).

Figure 2 shows the temperature dependence of the resource growth rate, foraging traits (attack rate and handling time), and metabolism for *C*_1_. The bell-shaped temperature response of the species attack rate (Fig. 2(b)) corresponds to the negative exponent of the Eqn. (3) and the U-shaped temperature dependence of handling time (Fig. 2(c)) corresponds to the positive exponent. We use parameter values that are feasible for ectothermic taxa (Sharpe and DeMichele 1977, Van der Have and De Jong 1996, Amarasekare and Coutinho 2014, Amarasekare 2015, Uszko et al 2017). Parameter values used in this study are given in the Tables 1 and 2.

**Table 1.** Model parameters, their definitions, and values. Parameters to fit into Boltzmann-Arrhenius relationship of the resource intrinsic growth rate and species metabolism, and unimodal Gaussian function of temperature response of consumer’s as well as predator’s foraging traits.

| Parameters | Ecological Interpretation | Value |
| --- | --- | --- |
| **Resource ($R$)** |  |  |
| $r_o$ | Intrinsic growth rate of resource at the reference temperature | 1 |
| $A_r$ | Arrhenius constant quantifying resource intrinsic growth temperature sensitivity | 10000 |
| $K$ | Strength of self-limitation | 6.7 |
| **Consumer ($C_1$)** |  |  |
| $a_{c_1 opt}$ | Maximal attack rate | 8.9 |
| $S_{a_{c_1}}$ | Performance Breadth for attack rate | 9.4 |
| $h_{c_1 opt}$ | Minimal handling time | 0.15 |
| $S_{h_{c_1}}$ | Performance breadth for handling time | 6.4 |
| $T_{opt a_{c_1}}$ | Temperature to attain maximum attack rate | 296K |
| $T_{opt h_{c_1}}$ | Temperature corresponding to minimal handling time | 294.1K |
| $e_{c_1}$ | Conversion efficiency | 0.3 |
| $m_{c_1 o}$ | Metabolism at the reference temperature $T_o$ | 0.25 |
| $A_{m_{c_1}}$ | Arrhenius constant quantifying temperature sensitivity of the species metabolism | 12000 |
| **Predator ($P$)** |  |  |
| $a_{p opt}$ | Maximal attack rate | 1.3 |
| $S_{a_p}$ | Performance Breadth for attack rate | 7.617 |
| $h_{p opt}$ | Minimal handling time | 0.1 |
| $S_{h_p}$ | Performance breadth for handling time | 11.25 |
| $T_{opt ap}$ | Temperature to attain maximum attack rate | 298K |
| $T_{opt h_p}$ | Temperature corresponding to minimal handling time | 298K |
| $e_{p_1}$ | Conversion efficiency | 0.7 |
| $m_{p o}$ | Metabolism at the reference temperature $T_o$ | 0.20 |
| $A_{m_p}$ | Arrhenius constant quantifying temperature sensitivity of the species metabolism | 12000 |
| $T_o$ | Ideal (reference) temperature | 297K |

**Table 2.**
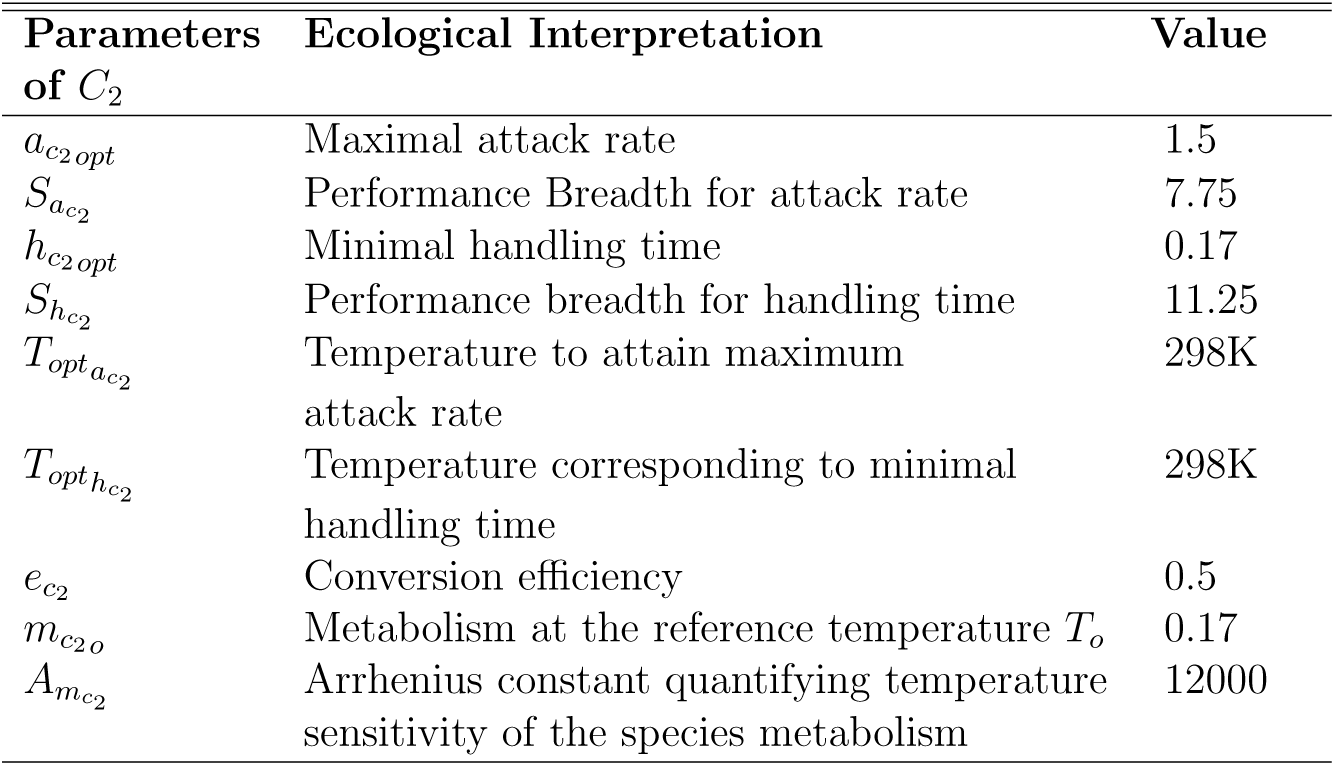
Additional parameters for the consumer *C*_2_ inhabiting in the diamond food web. Note that, parameters for all the other species (Eqn. (4)) are same as mentioned in the Table 1.

| Parameters of $C_2$ | Ecological Interpretation | Value |
| --- | --- | --- |
| $a_{c_2 opt}$ | Maximal attack rate | 1.5 |
| $S_{a_{c_2}}$ | Performance Breadth for attack rate | 7.75 |
| $h_{c_2 opt}$ | Minimal handling time | 0.17 |
| $S_{h_{c_2}}$ | Performance breadth for handling time | 11.25 |
| $T_{opt a_{c_2}}$ | Temperature to attain maximum attack rate | 298K |
| $T_{opt h_{c_2}}$ | Temperature corresponding to minimal handling time | 298K |
| $e_{c_2}$ | Conversion efficiency | 0.5 |
| $m_{c_2 o}$ | Metabolism at the reference temperature $T_o$ | 0.17 |
| $A_{m_{c_2}}$ | Arrhenius constant quantifying temperature sensitivity of the species metabolism | 12000 |

**Figure 2.**
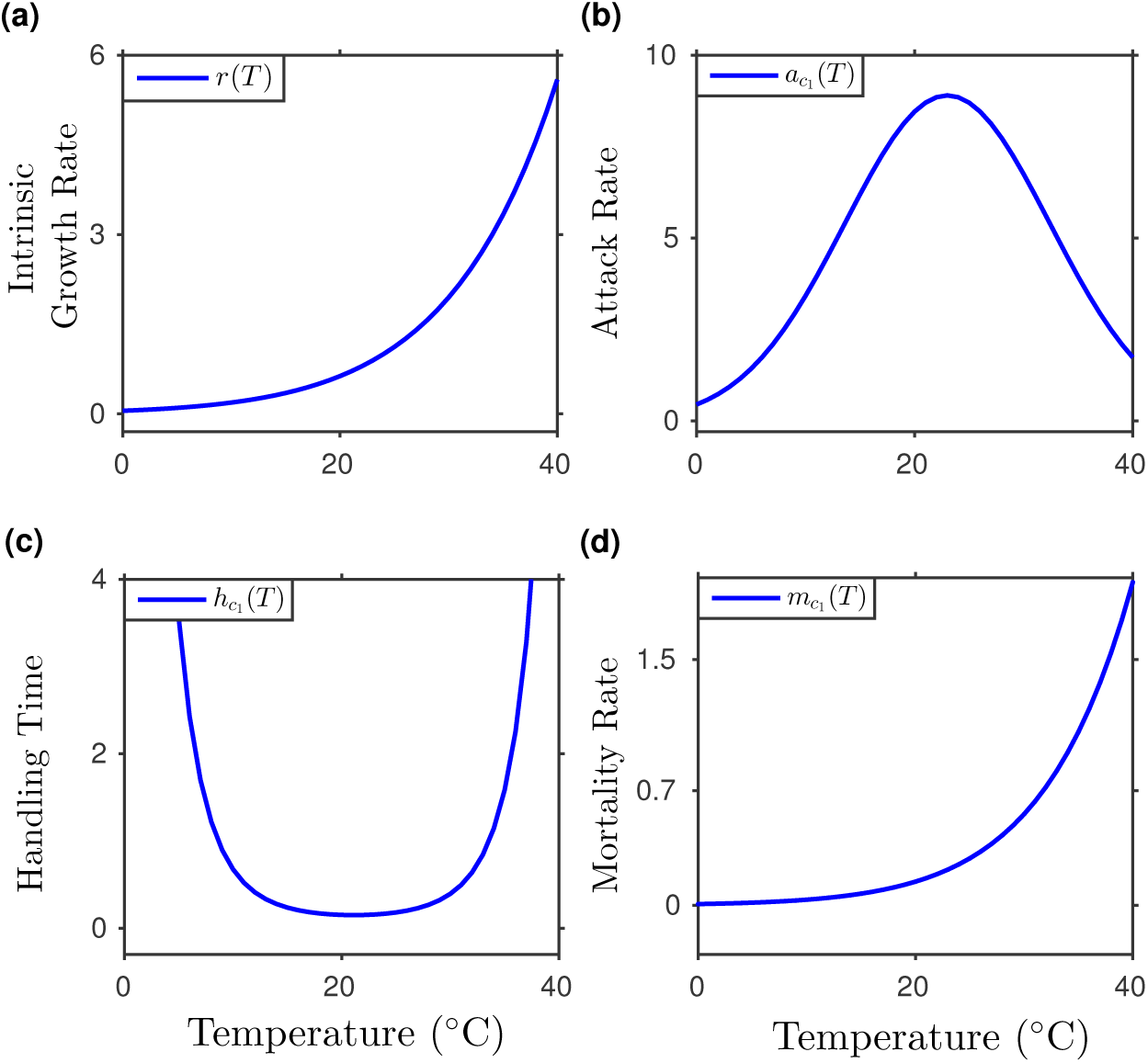
Monotonically increasing and unimodal temperature-dependent responses of species phenotypes. (a) Intrinsic growth rate *r* of *R* with variations in *T*. (b)-(c) Foraging capabilities of the consumer species, following the temperature dependence, where (b) is the bell-shaped thermal response of *C*_1_’s attack rate on *R*, and (c) is the U-shaped temperature-dependent behaviour of *C*_1_’s handling time. (d) Monotonically increasing temperature response of metabolic rate of *C*_1_. All the higher trophic species have a qualitatively similar temperature responses for their traits.

### Diamond food web

To understand how increasing the complexity influences ecosystem dynamics, we move from the food chain to investigate the diamond food web module (see Fig. 1(b)). The diamond food web is a four species food web module composed of energy pathways from a resource to two competitive consumers (apparent competition) vulnerable to predation by a top predator (Leibold 1996, McCann et al 1998, Levin 1970). We describe the food web model as:

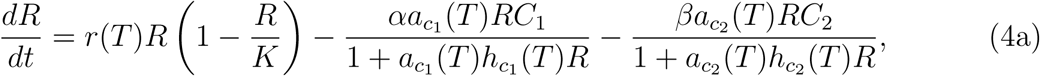

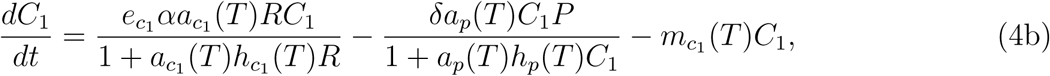

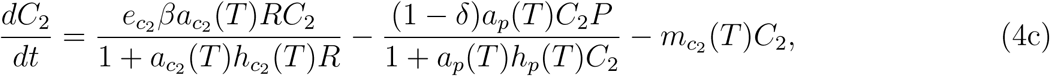

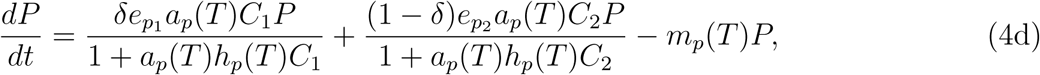

where *R* is consumed by two intermediate consumers; *C*_1_ and *C*_2_. Both *C*_1_ and *C*_2_ exhibit Type-II functional response over resource per-capita growth rate and are linked with *R* through preference parameters *α* and *β*, respectively. Further, the abundance of these consumers is under direct influence by *P* with preferences *δ* and 1 − *δ*, respectively. In Eqn. (4), 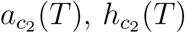 and 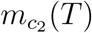 are temperature dependent physiological traits for consumer *C*_2_ having same definitions as described in Eqns. (2) and (3). 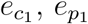 have same explanation as for previous models. While 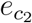 is conversion efficiency of *C*_2_ and 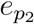 is the top predator’s conversion efficiency on the consumption of *C*_2_. Parameter values corresponding to biological traits of *C*_2_ are given in the Table 2.

### Omnivorous interaction

To elucidate the outcome of complex trophic interactions in response to warming, we increase the complexity further from the diamond food web to an omnivorous interaction (Fig. 1(c)). Omnivory is a feeding classification in which a species (an omnivore) consumes resources from more than one trophic level. The framework thus evolves from the food chain (Eqn. (1)) by incorporating an additional interaction between *R* and *P*, described as:

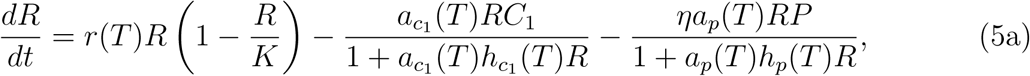

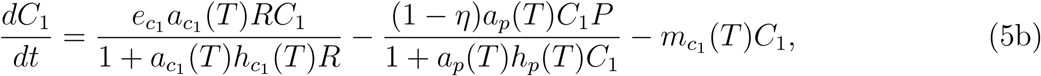

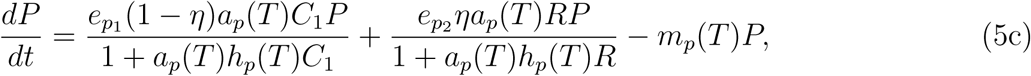

where the top predator *P* is a trophic omnivore. *P* can consume both the consumer *C*_1_ as well as the resource *R*. The parameter *η* governs the top predator’s preference for consumption of the two species, *R* and *C*_1_. Starting at *η* = 0, the system behaves the same as the tritrophic food chain (Eqn. (1)) with no omnivorous link. At *η* = 0.5, *P* has an unprejudiced preference; consumes both the resource and the intermediate species population in equal proportions (McCann and Hastings 1997). Any intermediate value of *η* determines the strength of omnivory. Increment in *η* value beyond 0.5, drags the system to exhibit intraguild predation where the predator is more of a competitor for the intermediate consumer (Hin et al 2011). Finally, at *η* = 1, the system transforms into a purely competitive form with two consumers (*P* and *C*_1_) competing with each other for the same shared resource (*R*). Here, 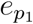 holds the same definition as described for Eqn. (1) and 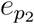 is the predator’s conversion efficiency on consuming *R*. The temperature dependence of the physiological traits is similar to that of the food chain and parameter values are described in the Table 1.

## Methods

We examine temperature dependence of species responses leading to their growth/consumption relative to the traits accounting for their metabolism or predation induced mortality, while moving from one trophic structure to another. Thus, analytical expressions characterising the relative responses for each of the *R, C*_*i*_ (*i* = 1, 2) and *P* depending upon their trophic interactions, are defined such that 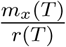 (*x* = *C*_1_, *C*_2_ or *P*, depending upon the trophic module) is the temperature-dependent metabolism of the species at higher trophic levels over the resource intrinsic growth rate. 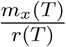 helps to track the metabolic requirements of species *x* relative to the available resource. Further, 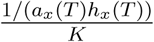 is the inverse enrichment ratio, which determines the resource density perceived by the species consuming the basal resource. Higher the value of 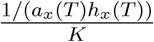 lower is the resource density consumed and vice-versa. Ω*/h*_*x*_(*T*) accounts for the maximum ingestion by species *x*, where Ω is the preference of *x* towards the ingested species. Thus, we define 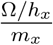 as maximum consumption by the species relative to their metabolism. We observe that in Eqns. (1), (4) and (5) species mortality is either due to energy lost during metabolism or due to predation. Therefore, species ingestion capabilities relative to their predation induced mortality is defined as 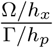, where Γ determines the preference of the top predator towards the consumer *x*.

Analyzing the thermal responses of these traits, first, we interpret species dynamics observed in each of the tri-trophic module for a specific choice of parameters. Later, for a comprehensive understanding of the realizable responses of the interaction modules towards warming, we perform the sensitivity of a few model parameters with temperature. It is well established that for the Type-II functional response, the resource carrying capacity *K* is a crucial metric for dampening or triggering oscillations in species density (Johnson and Amarasekare 2014, Amarasekare 2015). Initially, we study the system(s) sensitivity towards stability and persistence in the temperature–resource carrying capacity space, from one module to the other. Later, we investigate the robustness of interaction stability and persistence of species towards feasible alternative parameters: *α* and *β* (in the diamond food web), and omnivory strength *η* (in the omnivorous interaction).

Each of the deterministic systems is solved numerically either using MATLAB (R2015b) or the continuation package XPPAUT (Ermentrout 2002). For numerical integration, we use the 4th order Runge-Kutta method with adaptive step size.

## Results

Following the analytical expressions described in the methods section, qualitative behaviours of species relative physiological traits are depicted along the thermal gradient in Fig. 3. The increase or decrease in species abundance and extinction risk in the trophic structures due to changing temperatures are associated with the species relative traits.

**Fig. 3:**
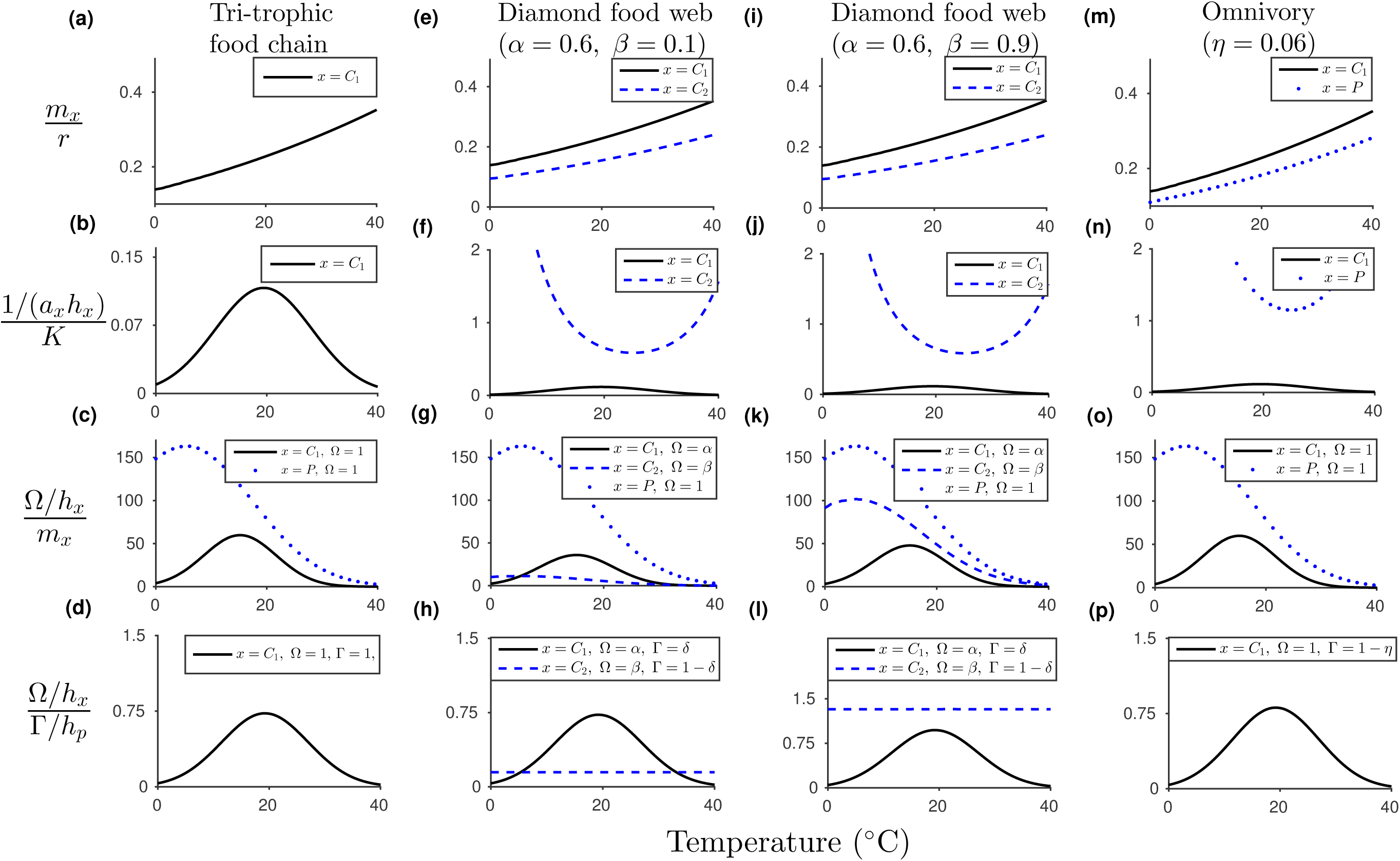
Thermal response of species physiological traits governing their growth relative to the metabolism or predation induced mortality. (a)-(d) Relative phenotypes for species in the tri-trophic food chain: (a) monotonically increasing behaviour of metabolism of 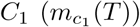 over resource growth rate (*r*(*T*)), (b) inverse measure of resource density perceived by *C*_1_, (c) temperature response of *C*_1_ and *P* ‘s maximum consumption relative to their metabolism (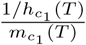 and 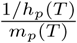, respectively), and (d) maximum consumption by *C*_1_ over its predation by *P*. (e)-(h) The behaviour of species relative foraging traits in accordance to the diamond food web module for the preference parameters *α* = 0.6 and *β* = 0.1: (e) metabolic rates of the competitors, *C*_1_ and *C*_2_ over the resource intrinsic growth rate (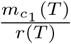 and 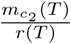 respectively), (f) inverse enrichmentratio of resource density as perceived by the consumers *C*_1_ and *C*_2_, (g) ingestion tendency of *C*_*i*_, *i* = 1, 2 over its metabolic rate, and (h) relative ingestion of the intermediate consumers *C*_*i*_ over their predation by *P*. (i)-(l) Similar relative traits of species in the diamond food web with preference parameters *α* = 0.6 and *β* = 0.9. (m)-(p) Temperature dependent relative behaviour of species traits interacting in the omnivorous model with omnivory strength *η* = 0.06: (m) metabolic rates of *C*_1_ and *P* over *r*(*T*), (n) thermal response of the resource abundance perceived explicitly by *C*_1_ and the omnivore *P*, (o) maximum ingestion of *C*_1_ and *P* over their respective metabolic rates, and (p) ingestion of *C*_1_ relative to *P* ’s predation rate.

### Dynamics of the tri-trophic food chain with increase in temperature

At very low temperatures, the food chain exhibits oscillatory dynamics via a Hopf bifurcation (HB), making the system unstable (see Fig. 4), while all the species co-exist. Further warming results in the stabilization of the oscillations via a reverse HB. We observe that increasing the mean temperature of the system (1) with a constant carrying capacity (*K*) increases 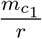 monotonically (see Fig. 3(a)). This indicates that resource growth rate increases relatively slower than the increase in the consumer’s metabolism. Thus, the metabolic requirements of *C*_1_ are higher than the growth of *R*. Furthermore, the bell shape of 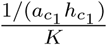 (Fig. 3(b)) reveals that the resource consumed by *C*_1_ decreases as the mean temperature increases from low to intermediate values. These factors reduce the flux of energy from the basal to the intermediate species. However, at the same time, maximum consumption of *C*_1_ increases at a faster rate than its metabolism as well as predation induced mortality (see Figs. 3(c)-3(d)). Thus, *C*_1_ is capable of surviving, but at a very low density. Meanwhile, the maximum consumption of the top predator increases faster than its metabolic requirements. The density of *P* increases at low temperatures, which further decreases gradually (see Fig. 4(c)) due to lesser availability of *C*_1_ as well as higher metabolic needs of the top predator with warming (see Fig. 3(c)).

**Figure 4.**
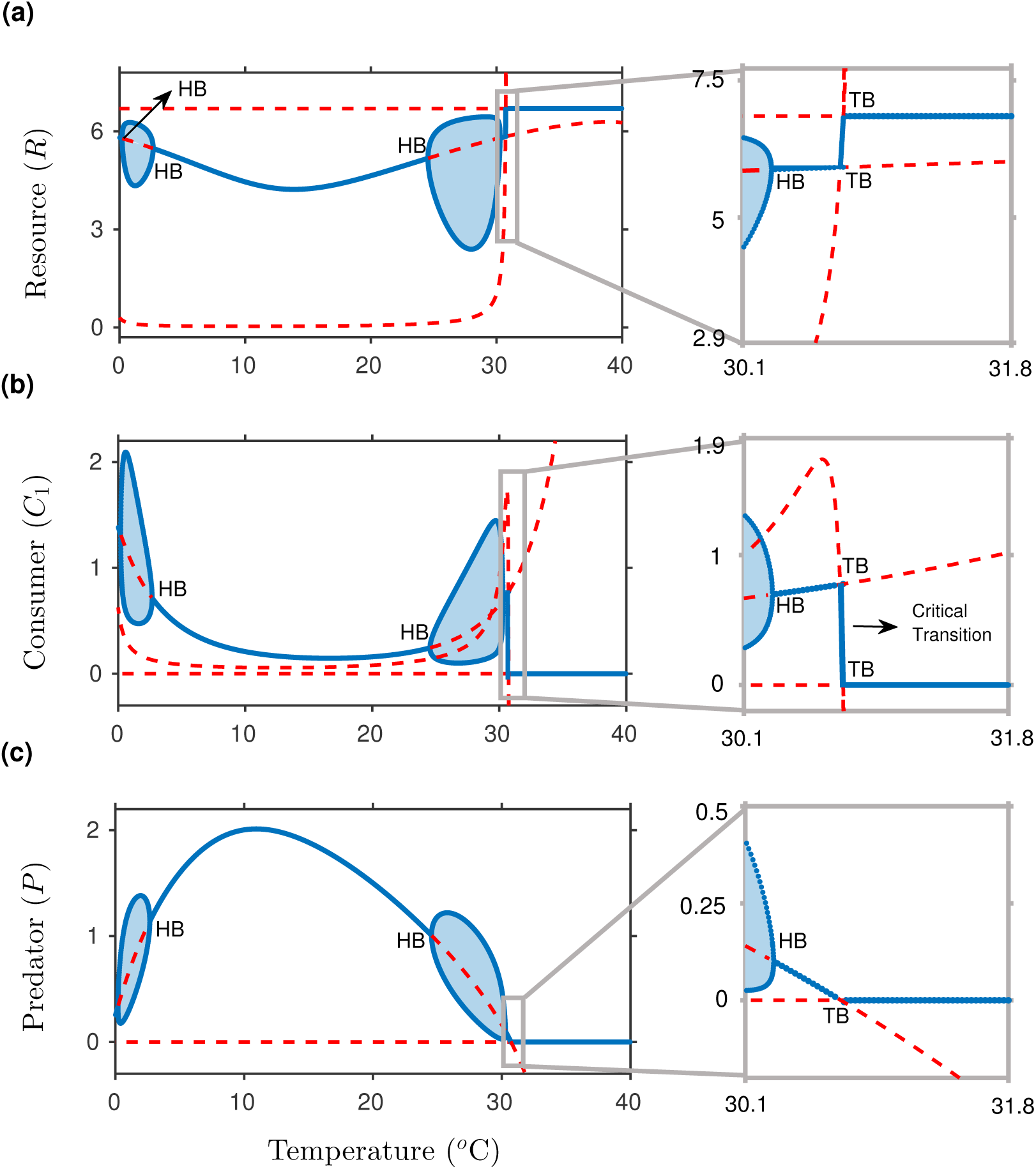
Species dynamics in the tri-trophic food chain (1) with variations in the temperature (*T*): for (a) the resource, (b) the consumer, and (c) the predator. The blow-up diagrams magnify the regions of species dynamics within the temperature range of 30.1°–31.8°C. The shaded regions describe the oscillatory state covering the upper as well as the lower amplitude of the species density. Solid lines (without shaded regions) determine the stable equilibrium densities, and dashed lines represent the unstable equilibrium densities. The Hopf bifurcation (HB) and the transcritical bifurcation (TB) represent a change in the qualitative behaviour of the species equilibrium. All the parameter values are the same as in the Table 1.

Within the intermediate temperatures (approximately between 15°–30°C), the system again bifurcates from the steady state to an oscillatory state via a HB and thus loses its stability (see Fig. 4). The proximity of low species density during oscillations is capable of driving them towards extinction. As the temperature crosses the intermediate range, the inverse enrichment ratio declines (see Fig. 3(b)), creating more availability of the resource for *C*_1_. However, on the other hand, the attack rate decreases, and the handling time increases for both the intermediate consumer and the top predator. As a consequence, species ingestion relative to their metabolism becomes very poor (see Figs. 3(c)). We notice a sudden collapse (with a tiny change in temperature) in the intermediate species density. As a result, both *C*_1_ and *P* become extinct at a high temperature via a transcritical bifurcation (TB) (shown in the blown-up diagram of Fig. 4(b)) and *R* reaches a steady state fixed at the carrying capacity *K*. Thus, the system shows stable dynamics but at the cost of the extinction of higher trophic species *C*_1_ and *P*.

### Dynamics of the diamond food web

In the diamond food web (Eqn. (4)), we investigate the impact of warming on the stability and the persistence of species for different interaction preferences, *α* and *β* of the competitors *C*_*i*_, *i* = 1, 2, respectively towards *R*. Our analysis starts with a weaker preference of the competitor *C*_2_ as compared to that of *C*_1_, towards consumption of *R*, and later we increase the interaction preference of *C*_2_. We also assume that the top predator has consumption preference (i.e. *δ* = 0.6) towards *C*_1_.

For weaker preference of *C*_2_ for *R*, i.e., *α* = 0.6 and *β* = 0.1, at low temperature range, the food web experiences a TB (see Figs. 5(a)-5(c)) and exhibits higher metabolic requirements by both *C*_1_ and *C*_2_, relative to *R*’s growth (see Fig. 3(e)). The combined consumption of the resource by the competitors reduces the resource density. Albeit resource abundance perceived by *C*_2_ is more than that by the competitor *C*_1_ (see Fig. 3(f)), it is observed that *C*_2_ is inferior to *C*_1_ in consuming the available resource (see Fig. 3(g)) relative to its metabolic rate. The selective up-take of the shared resource by the two competitors alters the energy flow within the interactive links in such a way that predation on *C*_2_ is higher than that on *C*_1_ (see Fig. 3(h)). Thus, low preference of *C*_2_ towards basal resource results in competitive exclusion (Gause 1934) of *C*_2_ by its competitor *C*_1_. Thus we observe that an interaction mediated by the top predator over the competitive consumers benefits the consumer *C*_1_ leading to the extinction of *C*_2_. Beyond the optimal temperatures for species traits, foraging capabilities of *C*_1_ are overpowered by its temperature-dependent mortality (see Figs. 3(g)-3(h)). We also observe that the addition of a competing species leads to early extinction of the higher trophic species, i.e., approximately at 29.2°C in comparison to that observed in the food chain (i.e., ≈ 30.7°C).

**Figure 5.**
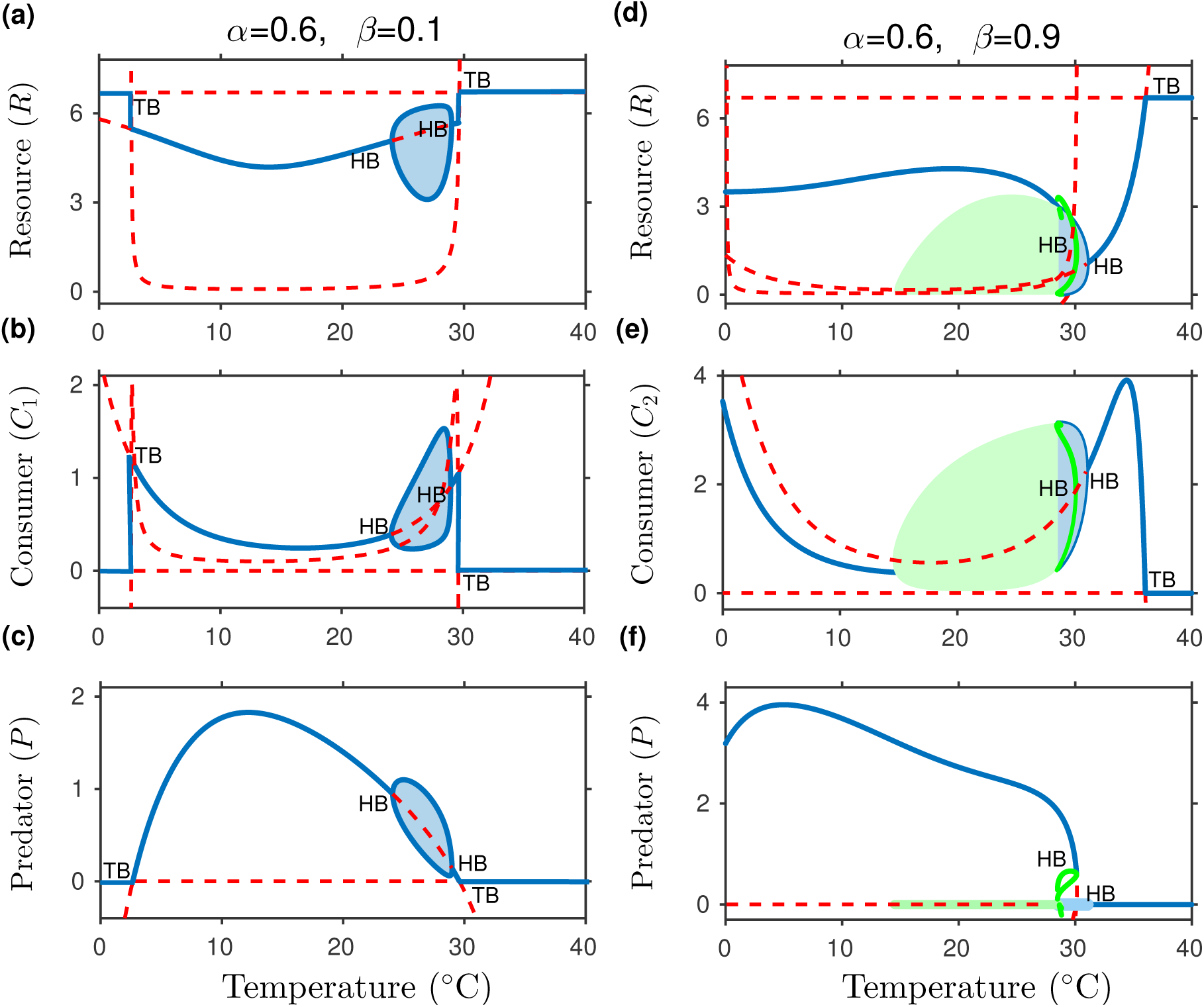
Species dynamics in the diamond food web along temperature gradient. (a)-(c) Bifurcation diagrams of the deterministic model having weak influence of the introduced competitor *C*_2_. Increasing the preference of the intermediate species (*C*_2_) for the basal resource, (d)-(f) depict the qualitative response of the surviving species towards warming. Green shaded regions and solid lines determine the existence of unstable limit cycles (i.e., the case when all the neighbouring trajectories approach it as time approaches negative infinity (Strogatz 1994)). Here, 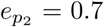 and the other parameter values are same as in Tables 1 and 2.

Following a stronger preference of *C*_2_ for *R* (i.e., *α* = 0.6 and *β* = 0.9), the major in- and fluence on species relative traits is visible on 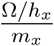 and 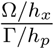 (see Figs. 3(k)-3(l)). At low temperatures, maximum consumption of the competitor *C*_2_ is higher than that of *C*_1_ (see Fig. 3(k)). Moreover, consumption capabilities of *C*_2_ relative to its predation induced mortality changes at same rate along thermal gradient (see Fig. 3(l)). In other words, ingestion abilities of *C*_2_ improves and overpowers the ingestion capability of *C*_1_. Higher preference of *C*_2_ and preference of *P* towards first consumer helps *C*_2_ to exclude *C*_1_ (see Figs. 5(d)-5(f)). Density of *C*_2_ declines and at the same time, there is an increase in abundance of *P* at intermediate temperatures. Further warming generates bi-stability exhibiting both, steady state (oscillation free) and oscillations. At *T* ≈ 28.6°C, we observe proximity of the predator density towards extinction (the predator density becomes very low), leaving negligible scope for the predator to escape extinction (see supplementary SI-2(a)). Particularly, *P* faces high risk of an early extinction in contrast to that of *C*_2_. However, both the higher trophic species *P* and *C*_2_ undergo mathematical extinction at *T* ≈ 36°C via a TB, mainly as a consequence of starvation due to their relatively higher metabolic rates poor foraging tendencies (see Figs. 3(k)-3(l)).

Overall, both the weak (*α* = 0.6 and *β* = 0.1) and the strong (*α* = 0.6 and *β* = 0.9) preferences of the additional consumer for shared resource exhibit competitive exclusion, leading to the extinction of one or the other intermediate consumer. Although the food web shows more stable dynamics than that of the food chain, the survival of the competing species depends upon their preference for the basal resource; species having a weaker preference towards *R* is expected to face extinction under the influence of temperature. Similar to the consequences in previous model, followed by the proximity of oscillatory dynamics towards extinction as well as the temperature dependence of species foraging, warming has an adverse effect on species persistence even at relatively low temperatures.

### Dynamics of the omnivorous interaction

Evolving from the linear flow of energy in the food chain module, here we consider the direct acquisition of the resource by the top predator (Eqn. (5)). We start our analysis by considering a weak influence of omnivory, where *P* has less effectiveness towards *R* and manifests preferential consumption of *C*_1_.

For *η* = 0.06, initial oscillations earlier observed in the food chain die out, making the omnivorous system more stable at low temperatures (see Figs. 6(a)-6(c)). Here, the relative physiological traits accounting for *C*_1_’s metabolic needs over *R*’s intrinsic growth (see Fig. 3(m)) and its ingestion capabilities over metabolism and density-dependent predation (see Figs. 3(o)-3(p)) are same as that observed in the food chain. The additional link between *P* and *R* reveals that *P* perceives resource abundance along with *C*_1_. The inverse enrichment ratio for *P* changes in a U-shaped manner (see Fig. 3(n)), indicating higher availability of the resource for the top predator at low to intermediate temperatures. At the same time 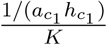 is similar for *C*_1_ (as observed in the food chain), indicating lesser availability of *R* for *C*_1_. Moreover, metabolic needs for *C*_1_ are higher than that of *P*. These factors reduce the flux of energy from *R* to *C*_1_, leading to consumer extinction at early temperatures and thus declining species co-existence. The top predator increases in its abundance (see Figs. 6(c) and 6(f)) due to the additional flow of energy through the resource.

**Figure 6.**
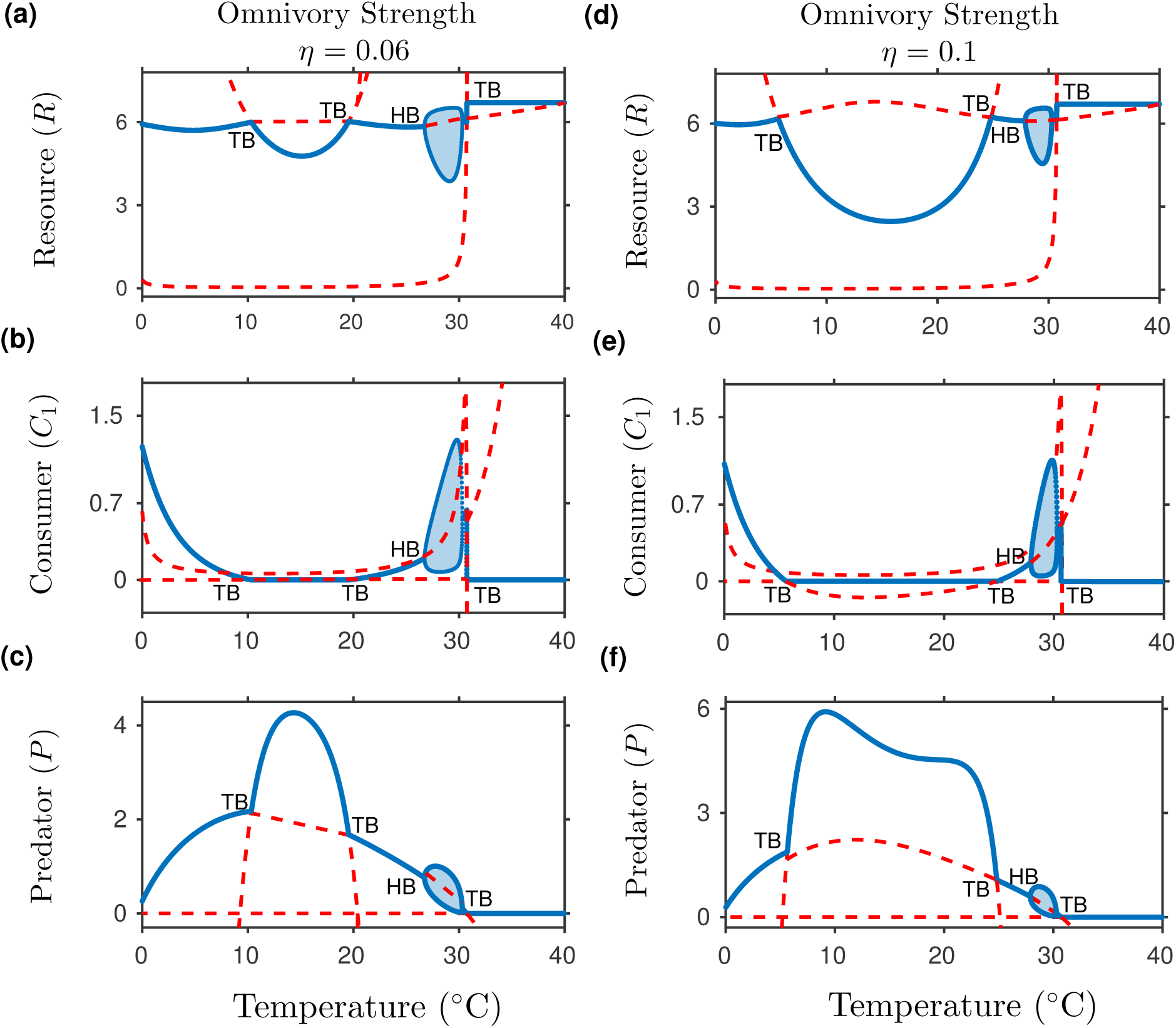
Variations in the species densities along temperature gradient for two different omnivory strengths *η* (see Eqn. (5)): (a)-(c) *η* = 0.06, and (d)-(f) *η* = 0.1. Model parameters are same as in the Table 1. Predator’s conversion efficiency for the resource intake is given by 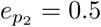.

Beyond the intermediate temperature range, delay in the HB threshold (now at an approximate temperature 26.7°C from 24.5°C in case of the food chain) reduces the temperature span of the non-equilibrium dynamics along with decreased amplitude of oscillations. Further, *C*_1_ re-invades the interaction network, but for a small temperature span (up to ≈ 30.3°C). Higher temperature (≈ 30.7°C) pushes the intermediate species and then the top predator to a consequence where their ingestion abilities cannot keep up with their metabolic rates (see Figs. 3(o)-3(p)). Thus, both the *C*_1_ and *P* become extinct due to starvation.

Increasing the omnivory strength to *η* = 0.1 further suppresses temperature span of oscillations in species density (27.9°C for *η* = 0.1 from 26.7°C for *η* = 0.06) (see Figs. 6(d)-6(f)). Beyond *η* = 0.14 and until *η* = 0.4, the intermediate oscillations become stable and the system shows stabilized dynamics throughout the temperature span (see supplementary SI-2(b)). However, when the omnivory strength is *η* = 0.5, the top most predator acts as a pure generalist with unbiased consumption of *R* as well as *C*_1_ (see supplementary SI-2(b)). The system changes its qualitative behaviour to unveil quasi-periodic oscillations, risking species survival due to proximity of lower amplitude of species density towards the extinction threshold (see Figs. 7(a)-7(c)). Overall, omnivory locally stabilizes an interior point equilibrium solution for a wide range of omnivory strength. Thus, increase in complexity from the food chain to the diamond food web and then to an omnivorous interaction suppresses oscillatory tendency of the system, however, it decreases the persistence of the intermediate consumer.

**Figure 7.**
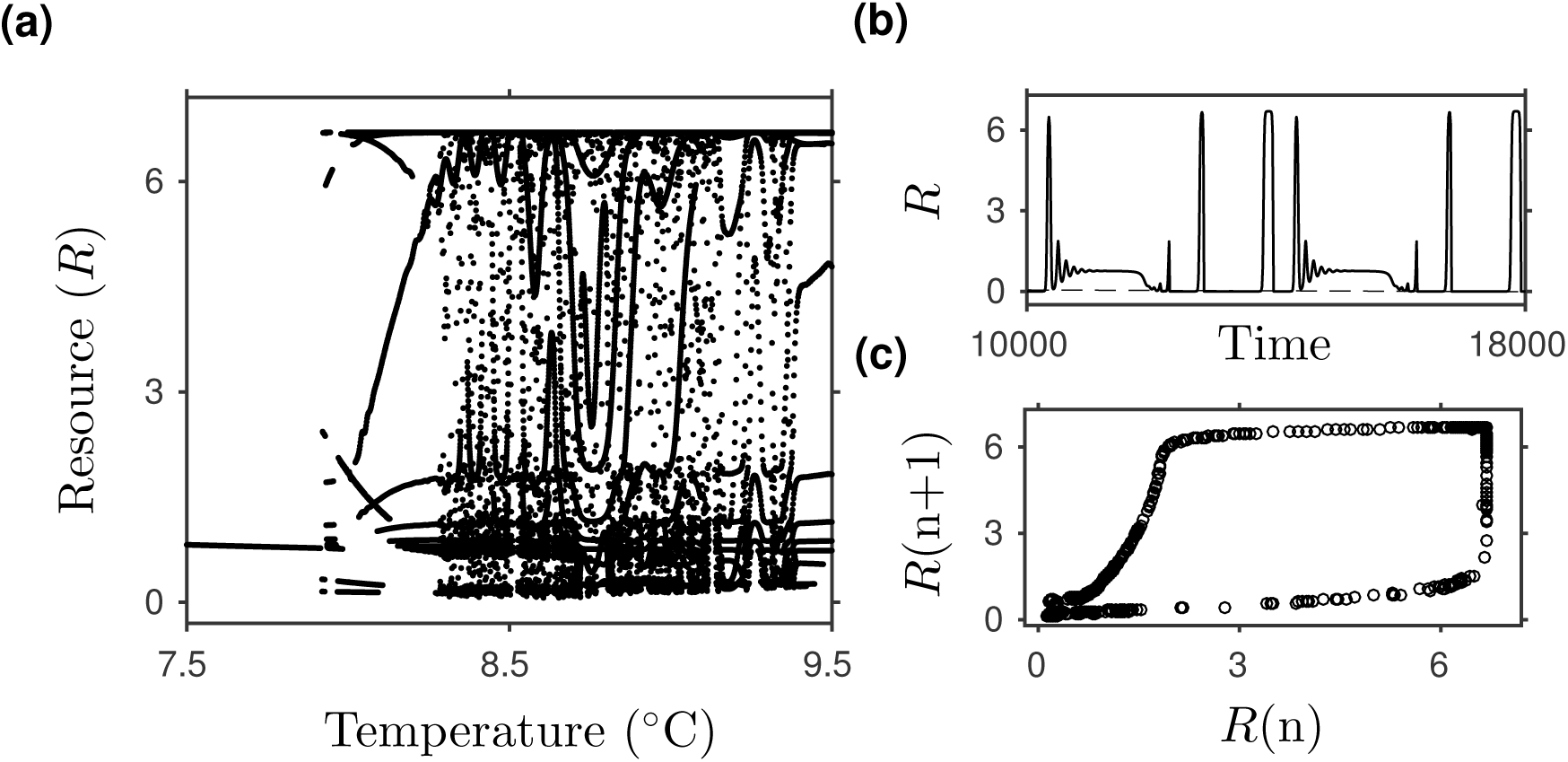
Species dynamics along the changing temperature in an omnivorous interaction: (a) resource dynamics for *η* = 0.5 exhibiting quasi-periodic oscillations, (b) time series obtained at *T* = 8.5°C depicting quasi-periodicity in the resource abundance, and (c) peak to peak plot obtained to track the behaviour of the ecological model at *η* = 0.5 and *T* = 8.5°C. Here, 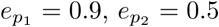 and 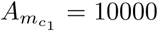. All the other parameters are same as in the Table 1.

### Sensitivity analysis

Up to now, the results presented are for fixed carrying capacity *K*, preference parameters *α* and *β* (for the food web), and omnivory strength *η* (for the omnivorous interaction), while the temperature is varied. To test the robustness of the dynamical outcomes observed in each of the tri-trophic modules, we perform sensitivity analysis in two-dimensional planes for a few experimentally accessible model parameters, through stability and persistence boundaries. The stability boundary corresponds to the HB curve (beyond which system exhibits oscillatory dynamics in species density), and the persistence boundary corresponds to the TB curve (beyond which species co-existing equilibrium fails to exist) (see Fig. 8).

**Figure 8.**
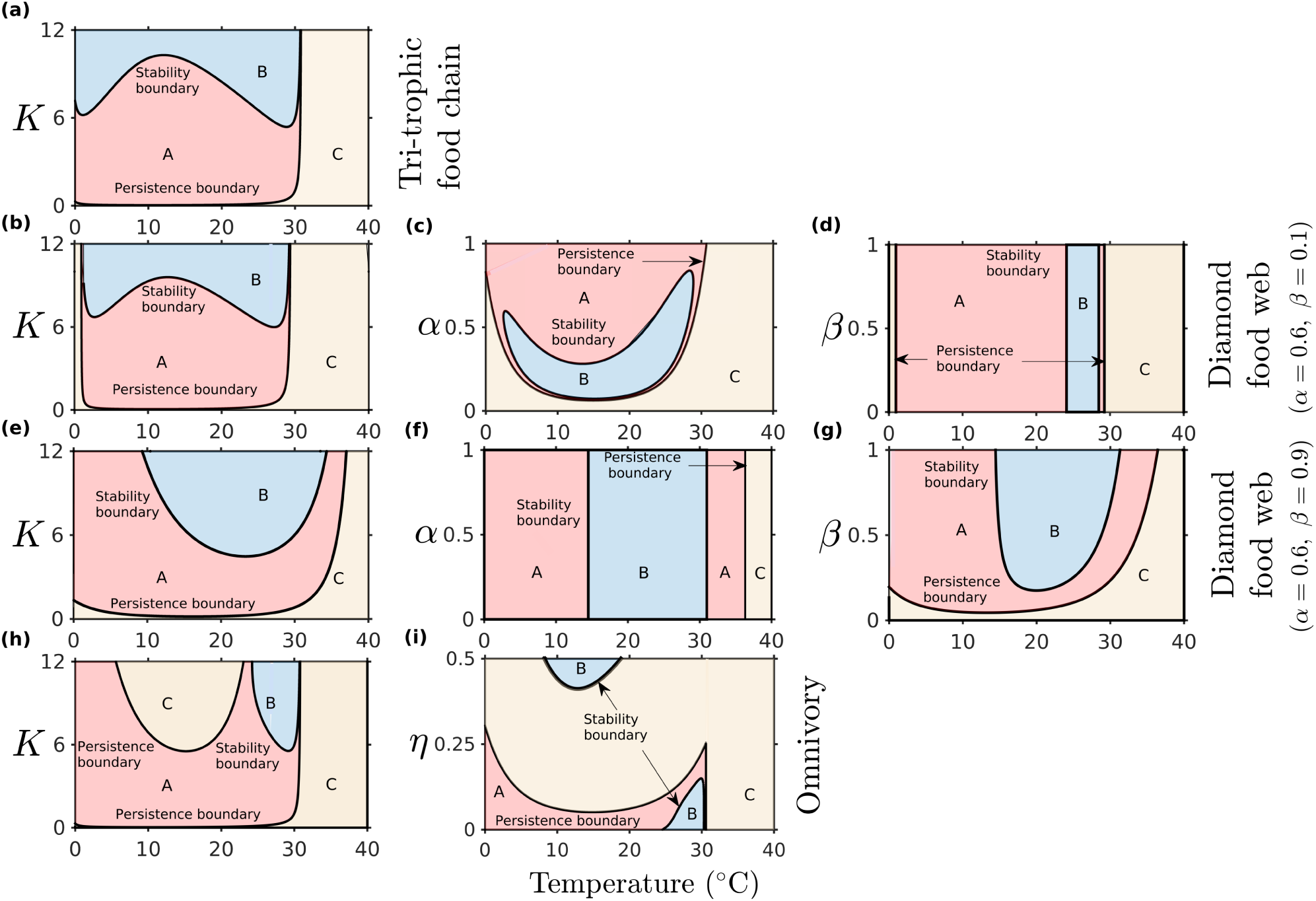
Species persistence and stability boundaries for each of the tri-trophic food chain, the diamond food web, and the omnivorous interaction in ((a), (b), (e), and (h)) temperature–resource carrying capacity space. Thermal sensitivity of preference parameters in the diamond food web for (c) *β* = 0.1, (d) *α* = 0.6, (f) *β* = 0.9, and (g) *α* = 0.6. (i) Sensitivity of the omnivory strength *η* towards systems stability and persistence along temperature gradient. Shaded regions: [A] determines steady state dynamics of the system(s), [B] is the region where the species density is associated with oscillatory behaviour, and [C] is the region beyond persistence boundary having no co-existence equilibrium.

It is visible that the species extinction is an inevitable outcome at high temperatures, in each of the trophic structures having low to high carrying capacity *K*. Further, moderate to high *K*, at low to intermediate temperatures, reveal oscillatory dynamics in the food chain (see Fig. 8(a)). In the diamond food web, the influence of resource carrying capacity towards the system’s stability depends upon the selective interaction of the competing consumers towards the basal resource (see Figs. 8(b)–8(g)). When *C*_2_ exhibits weak interaction preference in the diamond food web, the propensity of oscillatory dynamics with decreased persistence constrains within moderate to high carrying capacity as well as by the preference parameter of *C*_1_ for *R*. However, interaction preference of *C*_2_ with *R* does not influence the stability and persistence of species (see Fig. 8(d)). On the other hand, when the introduced competitor has stronger interaction preference with its resource, oscillations diminishes along the temperature axis and is constrained by the parameter *β* (see Figs. 8(e)-8(g)). Also, from the dynamical observations of Fig. 5(f), we note that in Figs. 8(e)–8(g), the steady state region [A] obtained crossing the right HB boundary at high temperatures indicates very low density of the predator, however, the mathematical extinctions of all the higher trophic species are realised on crossing the TB boundary. The initial oscillations further suppress in the omnivorous interaction, and species persistence drops at intermediate temperatures (see Fig. 8(h)). Therefore, as we increase the trophic complexity, the stability increases at the cost of the decrease in species persistence. Further, we observe that high temperatures are liable to exhibit oscillatory dynamics in the case of weak omnivory strength. Moderate to high *η* reveals a positive omnivory-stability relationship under the influence of temperature, though the co-existing equilibrium density of species fails to exist throughout the temperature axis (see Fig. 8(i)). Different conversion efficiencies of the consumer(s) and the predator yield similar outcomes along the temperature gradient (see supplementary SI-4). Overall, the sensitivity analysis depicts robustness of our observations concluding to increased stability and decreased persistence of species in complex interactions, along with changing temperature, for a wide range of model parameters.

## Discussion

In ecological interactions, the temperature–performance relationship serves as one of the essential factors to understand the impact of warming on biodiversity maintenance (Johnson and Amarasekare 2014). Significant alterations in community compositions and dynamics are evident from the interplay between species traits and temperature. Even with much progress, there remains a challenge in understanding the responses of trophic interactions towards changing temperature that links observations emerging through theoretical and experimental investigations. Thus, understanding the association of changing climatic conditions with well defined ecological interactions is necessary. Notably, it is essential to understand how the complexity of ecological interactions incorporating relevant performance-temperature relationship affects population dynamics. Here, we attempt to understand the impacts of warming on the persistence and stability of species having unimodal thermal response of their foraging traits.

We investigate species dynamics along the thermal gradient together with variations in their food web modules. We study temperature dependence of species biological traits to understand the dynamics of each of the tri-trophic modules. Our work concludes towards three main findings: First, by investigating three different tri-trophic food web modules, we find that species extinction is an inevitable outcome of warming, which is irrespective of their interaction network. Particularly, the extinction occurs due to the starvation as characterized by the declining temperature response of their foraging traits at high temperatures. Second, variations in the complexity of the considered food web modules can influence the effect of climate warming on species dynamics. A systematic increase in the complexity from the food chain to the omnivory reduces the propensity of oscillations in species dynamics. Thus, increasing trophic complexity in simple food webs can stabilize species dynamics, which is in accordance with few previous findings (MacArthur 1955, Elton 2000). Third, species co-existence drops on increasing the complexity, as the top-down control in the diamond food web fails, and warming does not support the survival of the intermediate consumer in the omnivorous interaction.

The above findings have key inferences for maintaining biodiversity. The first finding, indicating the extinction of species at high temperatures, emphasizes on the consequences of warming on community as well as individual species dynamics. It primarily focuses on the importance of considering the unimodal thermal response of species biological traits into models while investigating the ecological consequences of warming. Interestingly, it contrasts with the pattern of warming-induced extinction due to increased oscillations (Vasseur and McCann 2005). Since majority of the natural communities are composed of complex interactions, our other findings concern the detrimental effects of warming on more connected interactions in nature. It indicates that while larger communities may shrink due to species loss, simpler interactions exhibit more oscillations due to warming.

Recent theoretical and empirical evidences suggest that a large number of the Earth’s species had died out (Post 2013), and population declines signal the ongoing sixth mass extinction (Ceballos et al 2017). Using an experimental setup, Ullah et al (2018) demonstrated how warming leads to marine food web collapse through alterations in trophic flow. Here, while high temperatures influences the food chain to exhibit oscillation free behaviour, we also find that warming causes the extirpation of higher trophic species. Binzer et al (2012) incorporated monotonically increasing temperature dependence of species biological traits interacting in a food chain, and showed the extinction of the top predator, later followed by the loss of an intermediate consumer. Our work in contrast, studying the monotonically increasing growth and metabolic rates based on MTE and unimodal temperature responses of species foraging traits evident from empirical studies, eliminates any temperature delay in the extinction of the consumer as well as the top predator. Furthermore, the characteristic of ‘Paradox of Enrichment’ (Rosenzweig 1971) previously observed in bioenergetic consumer-resource model (Vasseur and McCann 2005), is also observed in the considered temperature dependent trophic structures for the increasing values of carrying capacity (see the left most panels in Fig. 8).

Increasing complexity of our framework by increasing species richness in the food chain, i.e., forming a diamond food web module, yields interesting dynamics along the temperature gradient. Literature proposes that varying metabolic demands of resources and consumers should intensify top-down control in consumer-resource systems (Levin 1970, Leibold 1996, Hoekman 2010, Shurin et al 2012, Dell et al 2014). In our work, keeping the basic structure of the food chain intact and gradually advancing the links within the species without altering the parametric values as well as temperature response of the species biological traits, we visualise a complete exclusion of one of the competitors, contrasting the top-down hypothesis. Křivan (2014) suggested that adaptive foraging by a predator in a diamond food web may yield dynamics similar to linear food chains. Here, we show that thermal dependence of species biological traits can also produce similar dynamics even by avoiding prey switching by the top predator. Thus, warming may create a stronger impact on species richness by reducing the impact of top-down control. Recently, Rudolf and Roman (2018) in an experimental set-up having interaction modules consisting of a resource consumed by two tadpole species (*Hyla versicolor* or *Rana clamitans*), exposed to a top predator (*Tramea carolina*) demonstrated the failure of the top-down control under ambient thermal conditions. They showed survival of the stronger competitor leading to the extinction of the weaker ones, and our results are also aligned with their findings. Our analysis further reveals that, in the diamond food web, the propensity to exhibit fluctuations in the species density decreases, however lower amplitudes of their densities are capable of pushing them towards extinction. The ecological system (Eqn. (4)) thus produces steady state dynamics but with increased risk of species extinction. Hence, moving from the food chain to the diamond food web, we observe the failure of species co-existence as well as increased vulnerability of intermediate consumer(s).

Furthermore, advancing from the food chain by increasing species connectance to the omnivory, we find that weak omnivory keeps the consumer density away from the extinction threshold at very low and high temperatures. However, an increase in the omnivory strength leads to the extinction of intermediate consumer throughout the temperature axis. At maximal omnivory strength, unstable dynamics due to increased oscillatory tendency of the species are witnessed within the intermediate temperature range (see Figs. 7(a)-7(b)). We generate a peak to peak plot to understand the dynamics occurring in the dense region of Fig. 7(a). The closed curve obtained in the peak to peak plot (see Fig. 7(c)) generated at 8.5°C represents the quasi-periodic behaviour of the omnivorous interaction. Earlier Pimm (1982) suggested that omnivory, when added to a system, is likely to make point attractors unstable, while our study indicates positive omnivory-stability relationship for moderate to high omnivory, under the influence of warming (see supplementary SI-2(b)). Beyond omnivory, Eqn. (5) demonstrates intraguild predation, where the predator has competitive dynamics with the consumer. Investigating thermal response of species abundance for intraguild predation reveals similar outcomes on species persistence and dynamical stability, indicating failure of species co-existence with warming (see supplementary SI-2(c)).

We consider the carrying capacity moderate enough to expect intrinsic oscillations by the interaction modules. As the carrying capacity can shape responses of the trophic structures towards warming, to check its sensitivity, the persistence and stability boundaries of the food web modules are plotted in temperature–carrying capacity plane (i.e., a two-parameter bifurcation diagram). In a large region of the parameter space, these boundaries have qualitatively similar responses as observed in the one-parameter bifurcation diagrams with variations in the temperature, i.e. increasing trophic complexity results in increased stability, but with higher propensity of species extinction, captured for moderate to high carrying capacity. Overall, our analysis holds significance across a range of ectothermic taxa assuming that higher trophic species metabolism increases faster than the resource intrinsic growth rate. Another assumptions requires a slower increase in species metabolic rates than their ingestion abilities at lower temperatures, followed by moderate to high temperature-independent resource carrying capacity.

To summarize, the rapid increase in the Earth’s temperature calls for a comprehensive understanding of its consequences on ecosystems. The study presented in this paper forms a basis that analyzes frameworks that can address questions of the effect of warming on the food webs, complexity-stability relationship, omnivory-stability relationship (May 1972, McCann 2000). We demonstrate the possibility of variable impacts of warming on different trophic interactions when considering declining performance-temperature relation at high temperatures. Earlier, May (1973) using Lotka-Volterra models found that trophic complexity is subject to destructive oscillations. Our results report contrary outcomes, however, under changing temperatures. Our work together with the temperature dependence of intraspecific competition may provide further insight to understand food web dynamics. Furthermore, climatic projections anticipate an increase in the intensity and frequency of temperature extremes. Also, the severity in declining species performance at high temperatures has a strong influence on temperature variation. Thus, work addressing periodic variability and stochastic variations in climatic warming in food webs may determine a wide range of effects of warming on trophic interactions.

## Acknowledgements

The authors acknowledge the Dutta group members for their suggestions on presentation. T.K. acknowledges Ramesh Arumugam for his help on computation.

