## SUPPORTING MATERIAL for "Persistence and stability of interacting species in response to climate warming: The role of trophic structure"

#### SI-1: Analytical expressions for species biological traits in response to changing mean temperature

We formulate temperature dependence of species biological traits defined in Eqns. (2) and (3) in the main text. Here, we express all the temperature dependent parameters explicitly corresponding to their Thermal Performance Curves (TPC's). Monotonically increasing thermal responses of the resource intrinsic growth rate ($r$), metabolic rates ($m_{c_i},\ i = 1,\ 2$) of the consumers ($C_1$ and $C_2$) and ($m_p$) of the top predator ($P$), respectively are given by:

$$r(T) = r_o\ e^{A_r(\frac{1}{T_o} - \frac{1}{T})}, \tag{SI-1.1}$$

$$m_{c_1}(T) = m_{c_1 o}\ e^{A_{m_{c1}}(\frac{1}{T_o} - \frac{1}{T})}, \tag{SI-1.2}$$

$$m_{c_2}(T) = m_{c_2 o}\ e^{A_{m_{c2}}(\frac{1}{T_o} - \frac{1}{T})}, \tag{SI-1.3}$$

$$m_p(T) = m_{p_o}\ e^{A_{m_p}(\frac{1}{T_o} - \frac{1}{T})}. \tag{SI-1.4}$$

Unimodal temperature dependence of the attack rates and the handling times of the consumers ($C_i,\ i = 1, 2$) and the predator $P$, respectively is given by:

$$a_{c_1}(T) = a_{c_1 opt}\ e^{-\frac{\left(T - T_{opt_{a_{c_1}}}\right)^2}{2S^2_{a_{c_1}}}}, \tag{SI-1.5}$$

$$h_{c_1}(T) = h_{c_1 opt}\ e^{+\frac{\left(T - T_{opt_{h_{c_1}}}\right)^2}{2S^2_{h_{c_1}}}}, \tag{SI-1.6}$$

$$a_{c_2}(T) = a_{c_2 opt}\ e^{-\frac{\left(T - T_{opt_{a_{c_2}}}\right)^2}{2S^2_{a_{c_2}}}}, \tag{SI-1.7}$$

$$h_{c_2}(T) = h_{c_2 opt}\ e^{+\frac{\left(T - T_{opt_{h_{c_2}}}\right)^2}{2S^2_{h_{c_2}}}}, \tag{SI-1.8}$$

$$a_p(T) = a_{p_{opt}}\ e^{-\frac{\left(T - T_{opt_{a_p}}\right)^2}{2S^2_{a_p}}}, \tag{SI-1.9}$$

$$h_p(T) = h_{p_{opt}}\ e^{+\frac{\left(T - T_{opt_{h_p}}\right)^2}{2S^2_{h_p}}}. \tag{SI-1.10}$$

Using the aforementioned analytical expressions, we represent physiological traits of $R$, $C_1$ and $P$ explicitly along the temperature axis (see Fig. 2 of the main text).

### SUPPORTING MATERIAL–2: From Taranjot Kaur & Partha Sharathi Dutta "Persistence and stability of interacting species in response to climate warming: The role of trophic structure"

##### SI-2: Species dynamics for different trophic structures

Here, we consider variations in the species preference parameters and investigate the dynamics of species abundance along with changing temperatures and trophic complexity.

*SI-2(a): Dynamics of the species in the diamond food web for $\alpha = 0.6$ and $\beta = 0.9$*

We observe that for high preference of $C_2$ towards $R$ (i.e. $\alpha = 0.6,\ \beta = 0.9$), species in the diamond food web exhibit bistable behaviour (see Figs. 5(d)–5(f) in main text). Moreover, the competitive (apparent) interaction leads to extinction of the intermediate consumer $C_1$ at low temperatures. The Fig. SI-2.1 displays magnified dynamics of species in the diamond

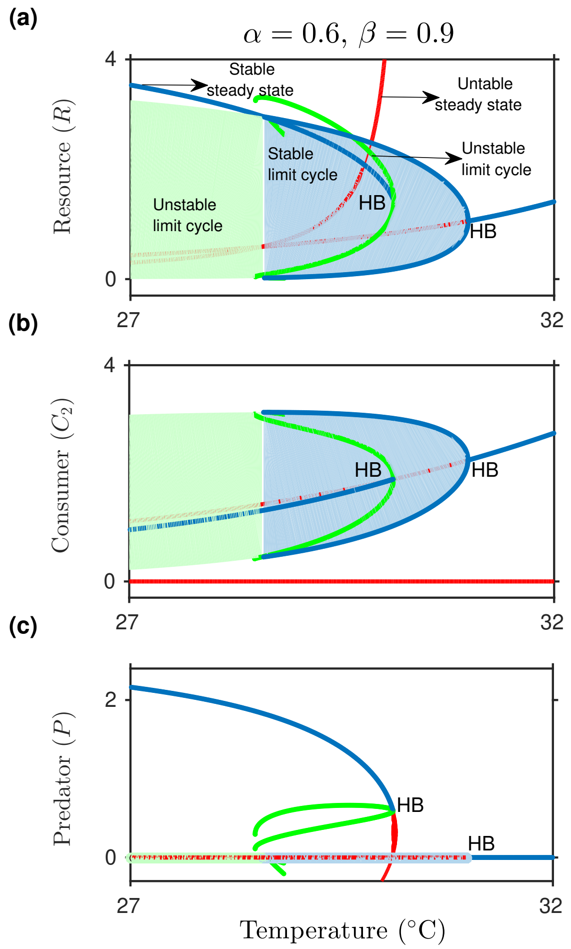


**Figure SI-2.1.** The dynamics of interacting species under influence of changing mean temperature $T$ from 27°C to 32°C for $\alpha = 0.6$ and $\beta = 0.9$. (a)-(c) Dynamics of $R$, intermediate consumer $C_2$ and $P$, respectively.

food web (see Figs. 5(d)-5(f)) for $\alpha = 0.6$ and $\beta = 0.9$. For the intermediate consumer $C_1$

the density hits the extinction at 0°C and stays there throughout the temperature gradient.

*SI-2(b): Omnivory-stability relationship*

Starting from low omnivory strength at $\eta = 0.06$ (see Fig. 6 in the main text) to high omnivory at $\eta = 0.4$, we analyse that the system exhibits stable dynamics while persistence of species with changing temperatures falls (see Fig. SI-2.2). Figure SI-2.2 represents time series of the species inherit in the omnivorous structure for different values of $\eta$. We observe that on increasing trophic complexity from the food chain to the omnivory, trade-off between the oscillations of $R$ as well $C_1$ tends to suppress the amplitude of oscillations in the top most predator's density (see Fig. SI-2.2(a)). Moving along the parameter $\eta$, oscillations die out from the system and initial dominance of $R$ is later suppressed by $P$ (see Figs. SI-2.2(a)-SI-2.2(d)).

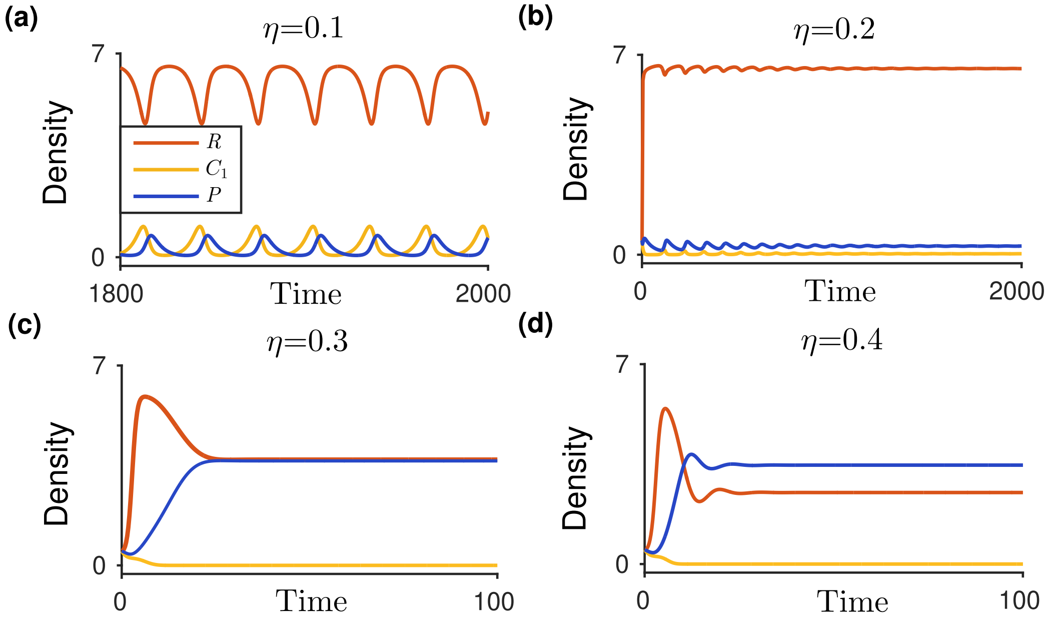


**Figure SI-2.2.** Time series representing species density for different values of omnivory strength $\eta$ at $T = 29.56$°C. (a) Oscillatory dynamics of the resource, the consumer and the predator at weak omnivory ($\eta = 0.1$), (b)-(d) completely stabilizing behaviour with increasing the omnivory strength from 0.2 to 0.4.

For $\eta = 0.5$, Here, we show complete dynamics of the system exhibiting maximum omnivory (i.e. $\eta = 0.5$). We find dense region of quasi-periodic oscillations (see Fig. 7

in the main text) in the system. The extinction risk of the species is higher in the dense region due to their proximity towards extinction boundary. On unveiling quasi-periodic oscillations, increase in the mean temperature pushes the system to exhibit steady state behaviour. Further, very high mean temperatures leads to starvation induced extinction of both $C_1$ and $P$, through transcritical bifurcation (TB).

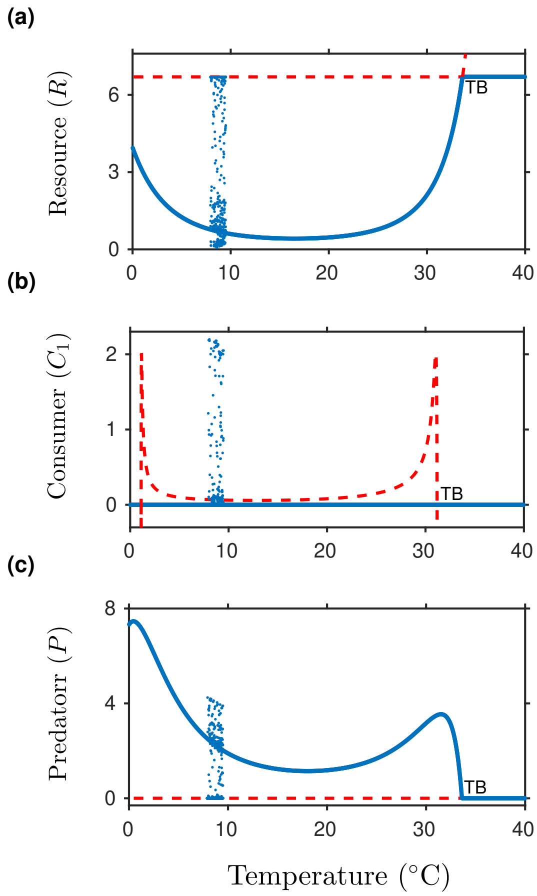


**Figure SI-2.3.** Species dynamics along the changing temperature when predator acts as a pure generalist ($\eta = 0.5$). Thermal response of the (a) resource (b) consumer, and (c) top predator. Dense region describes the quasi-periodic behaviour in species abundance.

*SI-2(c): Moving beyond omnivorous behaviour of the top predator*

Increasing $\eta$ beyond 0.5 pushes the ecological interaction expressed in the main text Eqn. (5) to a form of intraguild predation (McCann and Hastings 1997). Here, we analyse the thermal response of the basal resource $R$, the intermediate consumer $C_1$ as well as the top predator

$P$ for $\eta = 0.8$ and $\eta = 1.0$ (purely competitive model).

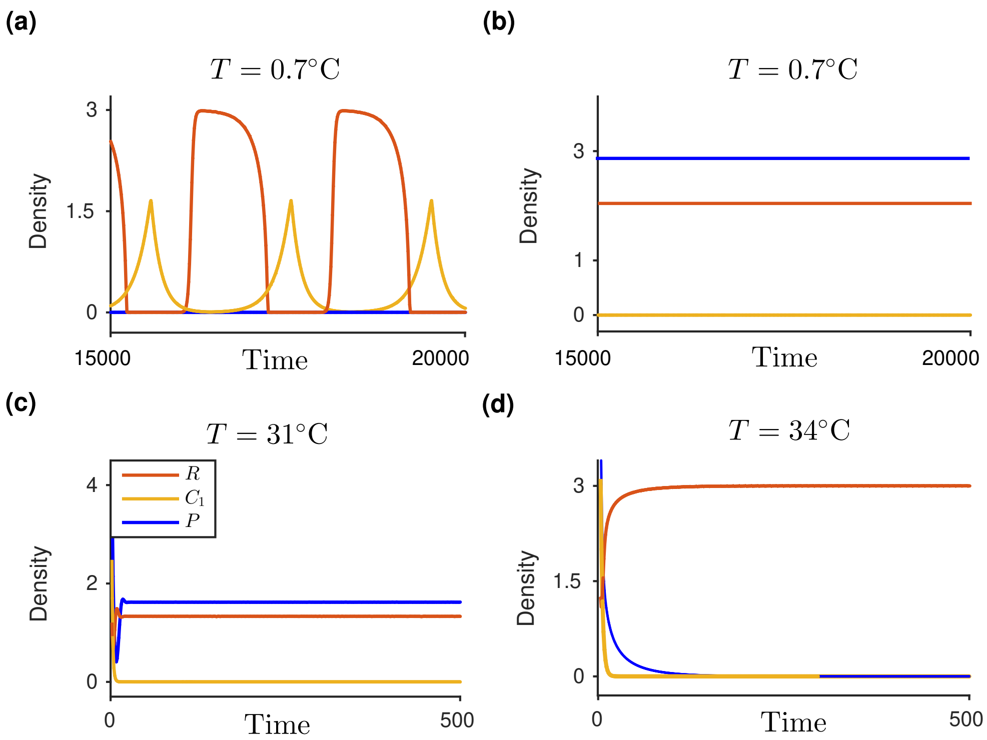


**Figure SI-2.4.** Time series analysis when the top predator is an intraguild predator. (a)-(b) Species density at $T = 0.7°C$ revealing bistable dynamics of the system at low temperatures. (c)-(d) Equilibrium abundance of the species and their proximity towards extinction for $K = 3.0$, $a_{p_{opt}} = 1.3$, $A_{m_p} = 12000$, $a_{c_1 opt} = 1.9$ and $A_{m_{c_1}} = 12000$. All the other parameter values are same as in the main text.

When predator is more of a competitor than an omnivore (i.e. $\eta = 0.8$), the time series analysis reveals that for low temperatures the interaction exhibits bistable behaviour showing both oscillatory as well as steady state dynamics (see Figs. SI-2.4(a)-SI-2.4(b)). Here, depending upon the initial densities, one of the intermediate consumer or the top predator undergoes warming induced extinction (see Figs. SI-2.4(a)-SI-2.4(b)).

As we increase the temperature beyond 12.5°C the system undergoes inverse Hopf bifurcation and oscillations die out leading to stabilized dynamics. The stabilized behaviour is liable to push the consumer and later the predator density towards extinction (see Figs. SI-2.4(c)-SI-2.4(d)). Thus, increasing the interaction complexity to an intraguild predation increases stability at the cost of higher trophic species extinction. For purely competitive behaviour of the top predator (i.e. $\eta = 1.0$) the interaction undergoes inverse Hopf bifurcation showing limit cycle induced oscillations at low temperatures (see Fig. SI-2.5). The inlet diagram in Fig. SI-2.5(b) depicts oscillatory dynamics in the phase plane for different values of temperature. We observe that these oscillations reveal very low predator density, leading to high risk of predator extinction at low temperatures. Moreover, risk towards species extinction is high even at intermediate temperatures. As the temperature increases, species density manifests stable behaviour but leads to extinction of $C_1$ later followed by $P$ via TB and $R$ reaches its carrying capacity. In all, moving interaction beyond omnivory is capable to catalyse species extinction along changing temperature.

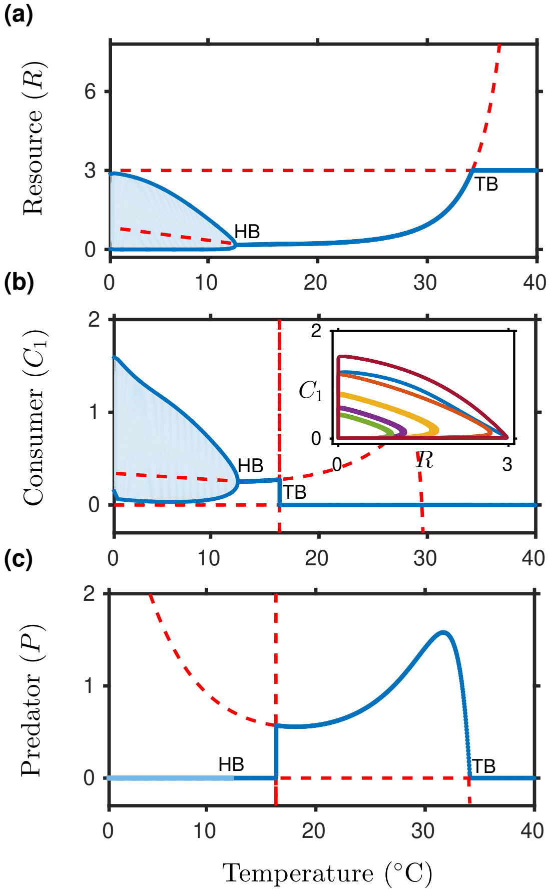


**Figure SI-2.5.** Qualitative response of species density towards warming in a pure competitive interaction for $\eta = 1.0$ and $K = 3.0$. (a)-(c) Competitive dynamics as well as local maximum and minimum density of $R$, $C_1$ and $P$, respectively. The inlet diagram depicts the phase plane analysis for temperature span within 0°C–12.5°C showing limit cycles of different amplitudes. Parameters are given by $a_{p_{opt}} = 1.3, A_{m_p} = 12000$, $a_{c_1 opt} = 1.9$, $A_{m_{c_1}} = 12000$ and other parameters are same as in the Table 1 (main text).

### SUPPORTING MATERIAL–3: From Taranjot Kaur & Partha Sharathi Dutta "Persistence and stability of interacting species in response to climate warming: The role of trophic structure"

#### SI-3: Unimodal temperature dependence of resource intrinsic growth rate

Studies on thermal response of species biological traits often reveal monotonically increasing or unimodal temperature dependence of the resource growth rate. Here, we extend our analysis as in the main text by incorporating $r(T)$ as a gaussian function of temperature such that

$$r(T) = r_{opt}\ e^{-\frac{(T-T_{opt_r})^2}{2S_r^2}}, \qquad \text{(SI-3.1.1)}$$

where $T_{opt_r}$ is temperature corresponding to the optimum resource growth rate $r_{opt}$ and $S_r$ is the temperature sensitivity of the resource growth. Considering unimodal thermal response of $r(T)$, we investigate the impact of changing mean temperature on stability and persistence of species in each of the tri-trophic modules studied in the main text.

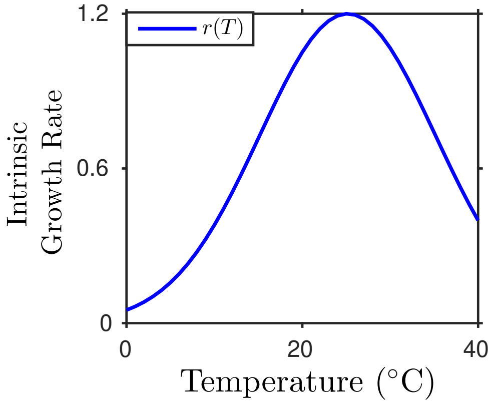


**Figure SI-3.1.** Unimodal temperature dependence of the resource intrinsic growth rate, followed from the Eqn. (SI-3.1.1).

##### *SI-3(a): Tri-trophic food chain*

The dynamics observed in the tri-trophic food chain for unimodal $r(T)$ are qualitatively similar to those observed when the resource growth rate increases monotonically along thermal gradient. At low temperatures, species density oscillates via a HB, spanning relatively (relative to that observed in main text) longer temperature range (see Fig. SI-3.2). Fur-

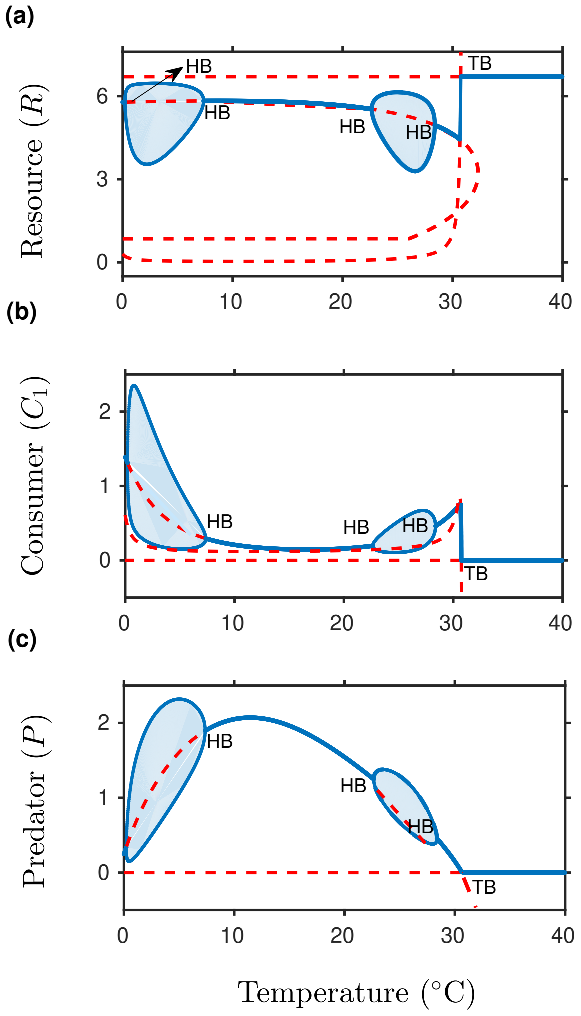


**Figure SI-3.2.** Species dynamics in the tri-trophic food chain along with changing mean temperature ($T$): for (a) $R$, (b)$C_1$, and (c) $P$. The resource growth rate follows unimodal temperature dependence (Eqn. (SI-3.1.1)) such that $r_{opt} = 1.2, T_{opt_r} = 298.15$K and $S_r = 10.0$. All the other parameter values are same as in the Table 1 of the main text.

ther warming leads to a reverse HB and shows steady state dynamics. We observe that at $\approx 30.7$°C (similar to that observed in the main text) the system stabilizes while the higher trophic species face extinction via a TB (see Figs. SI-3.2(b)-SI-3.2(c)). Moreover, before extinction of $C_1$ and $P$ at higher temperatures, all the species co-exist.

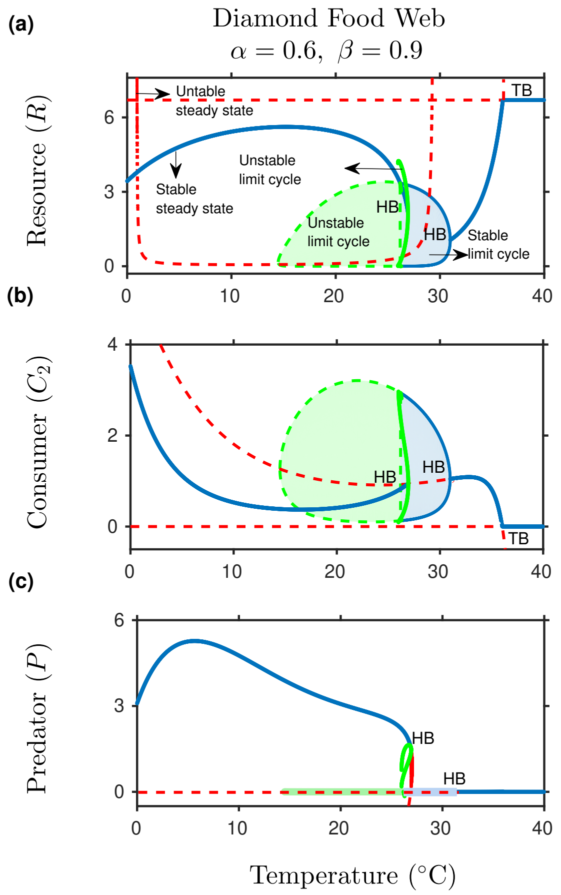


**Figure SI-3.3.** Temperature response of species interacting in the diamond food web. Steady state as well as oscillatory behaviour of (a) $R$, (b) the competitor, $C_2$ , and (c) $P$. The intermediate consumer $C_1$ hits extinction at 0°C and does not re-invade throughout the temperature range. All the parameters are same as in the Fig. SI-3.2.

#### *SI-3(b): Diamond food web*

We analyse system dynamics for $\alpha = 0.6$ and $\beta = 0.9$. At initial temperatures, species exhibit steady state dynamics. Increasing the temperature beyond optimal limits generates oscillatory dynamics (see Fig. SI-3.3). Furthermore, the unimodal temperature response of $r$ also fails to support top-down control and thus warming leads to extinction of the competitor $C_1$. Here, the system does not generate bi-stable states, though $P$ faces extinction at relatively lower temperatures than $C_2$ (see Figs. SI-3.3(b)-SI-3.3(c)). Thus, species co-existence is suppressed further than that in the food chain. At a very high temperature (≈ 30°C) $C_2$ also faces extinction and $R$ reaches its carrying capacity.

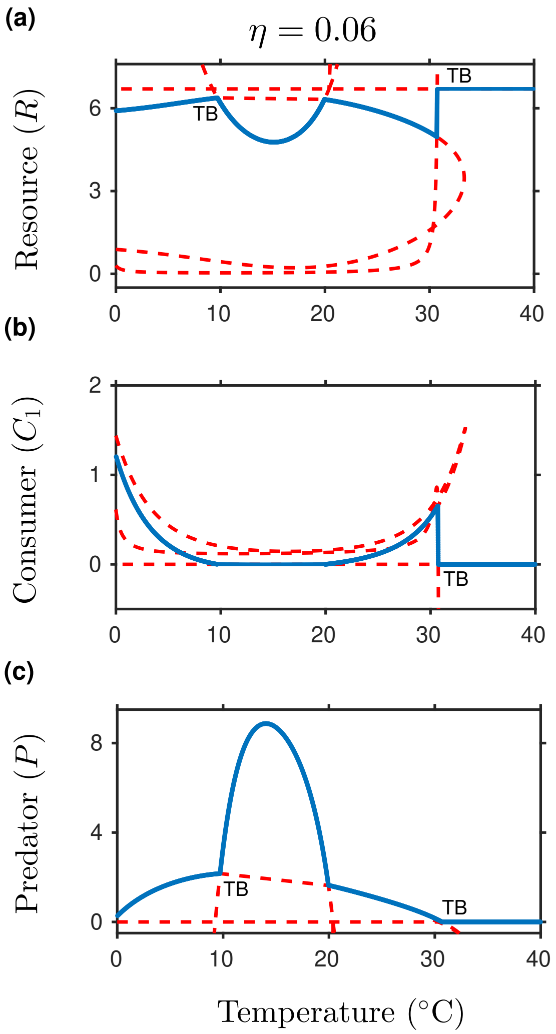


**Figure SI-3.4.** Species dynamics in the omnivorous interaction for $\eta = 0.06$, along the temperature axis: for (a) $R$, (b) $C_1$, and (c) $P$. All the parameter values are same as in the Fig. SI-3.2.

*SI-3(c): Omnivory*

We investigate the impact of warming on species persistence, moving from the food chain to the diamond food web and then to the omnivorous interaction. Here, we consider a weak omnivore i.e. $\eta = 0.06$.

On introducing an omnivorous link between the species in the food chain, we observe completely stabilized dynamics (see Fig. SI-3.4). The initial as well intermediate oscillations in the species abundance vanish. Moreover, species co-existence drops at low temperatures and $C_1$ cannot re-invade the system as observed when resource growth increases monotonically, along the thermal axis (see Fig. SI-3.4(b)). At a very high temperature i.e. $\approx 30.7°$C both $C_1$ and $P$ face extinction due to their poor foraging capabilities.

### SUPPORTING MATERIAL–4: From Taranjot Kaur & Partha Sharathi Dutta "Persistence and stability of interacting species in response to climate warming: The role of trophic structure"

#### SI-4: Temperature sensitivity of conversion efficiencies

Here, we investigate temperature sensitivity of conversion efficiencies ($e_{c_i},\ e_{p_i},\ i = 1, 2$) for each of the tri-trophic module studied in the main text.

Figure SI-4.1 depicts steady state, oscillatory as well as extinction regions of species for each module. In one hand, we observe that for the food chain a wide range of conversion efficiency of $C_1$ is liable to produce oscillatory dynamics in the system (see Fig. SI-4.1(a)). On the other hand, conversion efficiency of the top predator shows more sensitivity towards warming. For low values of $e_{p_1}$ (see Fig. SI-4.1(b)), the system shows complete stability from low to intermediate temperatures, followed by failure of co-existing species density at high temperatures. Increasing species diversity from the food chain to the diamond food web leads to similar observations, yet with stabilized dynamics and decreased persistence region (i.e. the region where all species co-exist) (see Figs. SI-4.1(c)–SI-4.1(j)). Furthermore, we observe that the omnivory increases stability, however with increased temperature span of the region having no co-existing equilibrium density (see Figs. SI-4.1(k)–SI-4.1(m)).

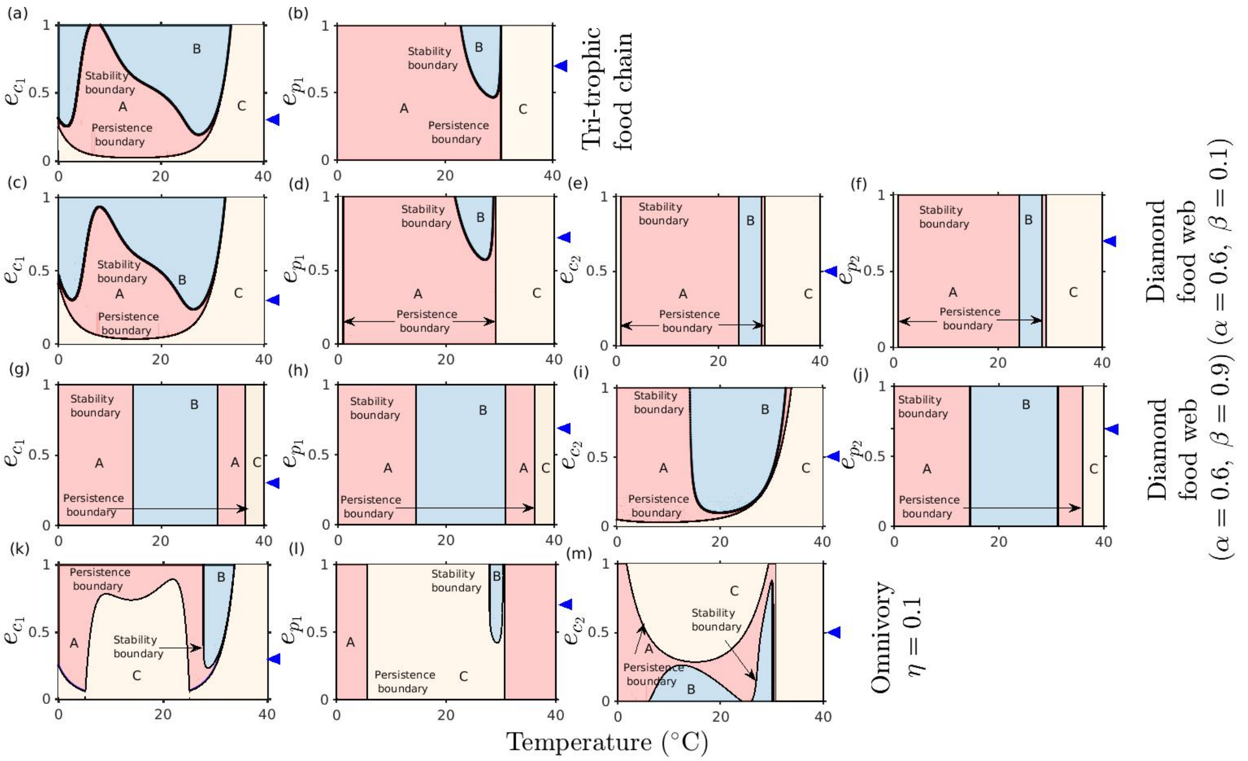


**Figure SI-4.1.** Thermal sensitivity of species conversion efficiencies for the: (a)-(b) food chain, (c)-(j) diamond food web (in (c)-(f) $\alpha = 0.6$, $\beta = 0.1$, and (g)-(j) $\alpha = 0.6$, $\beta = 0.9$), and (k)-(m) omnivorous interaction ($\eta = 0.1$). Shaded regions: [A] determines steady state dynamics of the system(s), [B] is the region where the species density is associated with oscillatory behaviour, and [C] is the region beyond persistence boundary having no co-existence equilibrium. The blue triangles mark the parameter values used in the main text to calculate the bifurcation diagrams for each module.
